# TREM2 impacts brain microglia, oligodendrocytes and endothelial co-expression modules revealing genes and pathways important in Alzheimer’s disease

**DOI:** 10.1101/2021.07.16.452732

**Authors:** Guillermo Carbajosa, Karim Malki, Nathan Lawless, Hong Wang, John W. Ryder, Eva Wozniak, Kristie Wood, Charles A. Mein, Alan Hodgkinson, Richard J.B. Dobson, David A. Collier, Michael J. O’Neill, Stephen J. Newhouse, Angela K. Hodges

## Abstract

A microglia response to pathogenic signals in diseases such as Alzheimer’s disease (AD) has long been recognised, but recent genetic findings have cemented their direct causal contribution to AD and thus the potential to target them or their effector pathways as a possible treatment strategy. TREM2 is a highly penetrant microglia risk gene for AD, which appears central to the coordination of the damage response by microglia in AD. Its absence has a negative impact on Tau and amyloid symptoms and pathologies. Full knowledge of its pathway and relationships with other brain cells in AD has not been fully characterised, but will be essential to fully evaluate the impact of manipulating this pathway for treatment development and to establish the best targets for achieving this. We used whole genome RNA sequencing of hippocampus and cortical brain samples from control, AD, and AD TREM2 risk carriers to identify TREM2-dependent genes driving changes in pathways, processes and cell types in AD. Through highly influential intra and intermodular hub genes and overall changes in the levels of gene expression, TREM2-DAP12 was found to strongly influence a number of other microglia, oligodendrocyte and endothelial genes, notably those involved in complement and Fcγ receptor function, microglia-associated ribosomal genes and oligodendrocyte genes, particularly proteosomal subunits. These strong TREM2 centred co-expression relationships were significantly disrupted in AD cases with a TREM2 risk variant, revealing for the first time genes and pathways directly impacted by TREM2 in the brains of AD patients. Consistent with its function as a lipid sensor, our data supports a role for TREM2 in mediating oligodendrocyte and/or myelin clearance in AD which may be essential not only for preserving healthy tissue homeostasis but may also serve to minimise the persistence of antigenic peptides and lipids which may lead to detrimental pro-inflammatory sequelae. Further work should expand our knowledge of TREM2 on complement and Fcγ receptor function and its impact on oligodendcrotye and myelin integrity and further evaluate the genes and pathways we have identified as possible treatment targets for AD.

## Introduction

Rare missense variants in TREM2 (1-2% frequency in AD) can significantly increase the risk of developing Alzheimer’s disease (AD) and related neurodegenerative diseases (Guerreiro *et al*., 2013b; Guerreiro *et al*., 2013c; Jonsson *et al*., 2013; Jin *et al*., 2014; Ridge *et al*., 2016). As TREM2 in the brain appears to be exclusively expressed in microglia, this provides compelling evidence that microglia are involved in vulnerability to neurodegeneration. TREM2 is a transmembrane receptor containing an extracellular immunoglobulin V-set domain suggesting potential for high and low avidity interactions with a broad range of ligands reported to include Aβ (Lessard *et al*., 2018; Zhao *et al*., 2018), apolipoproteins (Atagi *et al*., 2015; Yeh *et al*., 2016), anionic and zwitterionic lipids (Wang *et al*., 2015) and anioinic carbohydrates (Daws *et al*., 2003).

Individual variants are believed to impact TREM2 production (Q33X), expression or turnover at the cell surface (R47H) (Kleinberger *et al*., 2014; Park *et al*., 2017), α-secretase cleavage of an extracellular soluble ectodomain (H157Y) (Kleinberger *et al*., 2014; Jiang *et al*., 2016a; Thornton *et al*., 2017) and/or ligand binding (R47H, R62H, T96K, H157Y) (Kober *et al*., 2016; Song *et al*., 2017; Sudom *et al*., 2018) leading to loss of normal TREM2 function. Additional rare missense variants affecting both alleles primarily alter protein structure, cell surface expression and shedding, and cause Nasu-Hakola disease (Paloneva *et al*., 2002; Kleinberger *et al*., 2014; Kober *et al*., 2016). This very rare fatal disease emerging in early adulthood is characterised by fragility bone fractures, demyelination, calcification of the basal ganglia and progressive frontal dementia (Paloneva *et al*., 2001; Bock *et al*., 2013). There are clear overlaps with AD, although Nasu-Hakola patients fail to display appreciable levels of amyloid or Tau deposits in the brain (Satoh *et al*., 2018). These evidence serve to highlight the pleiotropic nature of TREM2 activity and highlight the relevance of temporal and spatial context. In fact, there are widely differing ages of disease onset, disease symptoms and pathology in the TREM2 AD patients we have evaluated, even those with the same TREM2 risk variant. Disease manifests just like any other patient with AD, FTD or similar condition, making TREM2 carriers indistinguishable from other patients in the absence of genetic knowledge. This suggests partial loss of TREM2 function by itself is insufficient for disease and that a number of additional stochastic factors can greatly influence disease risk. It also suggests that dysfunctional TREM2 linked pathways in microglia may be central to AD pathogenesis even in patients without a predisposing genetic risk in TREM2.

Over-expression or restoration of normal TREM2 function has been shown to have largely beneficial outcomes in amyloid-driven (Jiang *et al*., 2014; Lee *et al*., 2018; Song *et al*., 2018) or Tau-driven AD mouse models (Jiang *et al*., 2016b). This suggests that the microglia response observed in AD brain might at least in part be an attempt, albeit one that ultimately fails in AD patients, to protect from AD pathogenesis. Loss of normal TREM2 function via knock-out or introduction of a disease variant suports this. In an amyloid background, loss of TREM2 leads to an exacerbation of AD pathology. There are fewer microglia, defective production of soluble TREM2 ectodomain, less microglia clustering around plaques (Song *et al*., 2018), impaired plaque barrier function and fibrillar Aβ, plaque and plaque-associated APOE clearance (Wang *et al*., 2015; Yuan *et al*., 2016; Parhizkar *et al*., 2019). Behavioural data to support these findings is however currently lacking and some studies have reported lower plaque load in the absence of TREM2 (Jay *et al*., 2015; Krasemann *et al*., 2017). Tau models lacking TREM2 further support the protective function of normal TREM2, displaying exacerbated Tau pathology, more ramified and fewer microglia, increased brain atrophy and greater memory deficits (Jiang *et al*., 2015; Bemiller *et al*., 2017; Sayed *et al*., 2018). Somewhat at odds with this, Leyns *et al*. (2017) and Sayed *et al*., (2018) note that lower cytokine production, less atrophy and synaptic loss occurs in Tau mice completely lacking TREM2 (Leyns *et al*., 2017; Sayed *et al*., 2018), suggesting TREM2 may respond differently depending on the timing of microglia exposure to pathological cues such as amyloid, aggregated Tau or neuronal cell loss which emerge at different ages and stages of disease progression. It is notable that the PS19 Tau line (Leyns *et al*., 2017; Sayed *et al*., 2018) has greater and earlier neuronal cell loss, brain atrophy and microgliosis than the hTau line (Bemiller *et al*., 2017) and haplosufficiency versus complete loss of TREM2 may also be a factor. Loss of TREM2 has also been shown to lead to deficits in phagocytosis, at least in the uptake of Aβ, myelin debris, apoptotic neurons and/or *E.coli* (Takahashi *et al*., 2005; Kleinberger *et al*., 2014; Cantoni *et al*., 2015; Xiang *et al*., 2016; Garcia-Reitboeck *et al*., 2018). These effects are consistent with the known signalling pathways linking TREM2, through it’s adaptor DAP12 and P13K, PLCγ and VAV/RAS/ERK which impact cell survial, proliferation, cytokine and chemokine secretion and phagocytosis, reviewed in (Ulland and Colonna, 2018).

Expression profiling of mouse and human tissue has significantly increased our knowledge of markers specific to different brain cell types and revealed important cellular subpopulations with unique molecular identities and behaviours in normal (Hawrylycz *et al*., 2012; Forabosco *et al*., 2013; McKenzie *et al*., 2018; Olah *et al*., 2018) and AD brain (Zhang *et al*., 2013). Recent single-cell RNA sequencing studies have further refined important molecular profiles representing different microglia cell sub populations in human brain (Mathys *et al*., 2017; Del-Aguila *et al*., 2019; Patir *et al*., 2019; Sankowski *et al*., 2019; Srinivasan *et al*., 2019) some of which (DAM, damage-associated microglia) co-localise to disease-relevant structures such as amyloid in mouse amyloid models of AD (Keren-Shaul *et al*., 2017) or associate with myelinating oligodendrocytes (PAM, proliferative-region-associated microglia) during normal early development (Li *et al*., 2019). They also highlight the central position of TREM2 and TREM2-associated networks within specific microglia subpopulations and their likely rippling impacts on other brain cells types associated with AD (Carbajosa *et al*., 2018).

We therefore sought to identify genes, processes and cell type changes driven specifically by TREM2 in human AD by comparing expression profiles in control, and AD patients with and without an AD-associated TREM2 variant. Modules containing genes with highly correlated patterns of gene expression and enriched for different cell identities revealed interactions and highly influential intra and intermodular hub genes. They also allowed us to establish TREM2- dependent and independent pathways impacting AD and establish which cell types are being impacted in AD by TREM2. The goal was to establish the influence of microglia within the whole brain AD environment in people, identify molecular pathways linked to TREM2 and establish a benchmark by which TREM2-related mouse models can be evaluated.

## Methods

An overview of the data analysis pipeline used in the study can be found in Figure 1.

**Figure 1:**
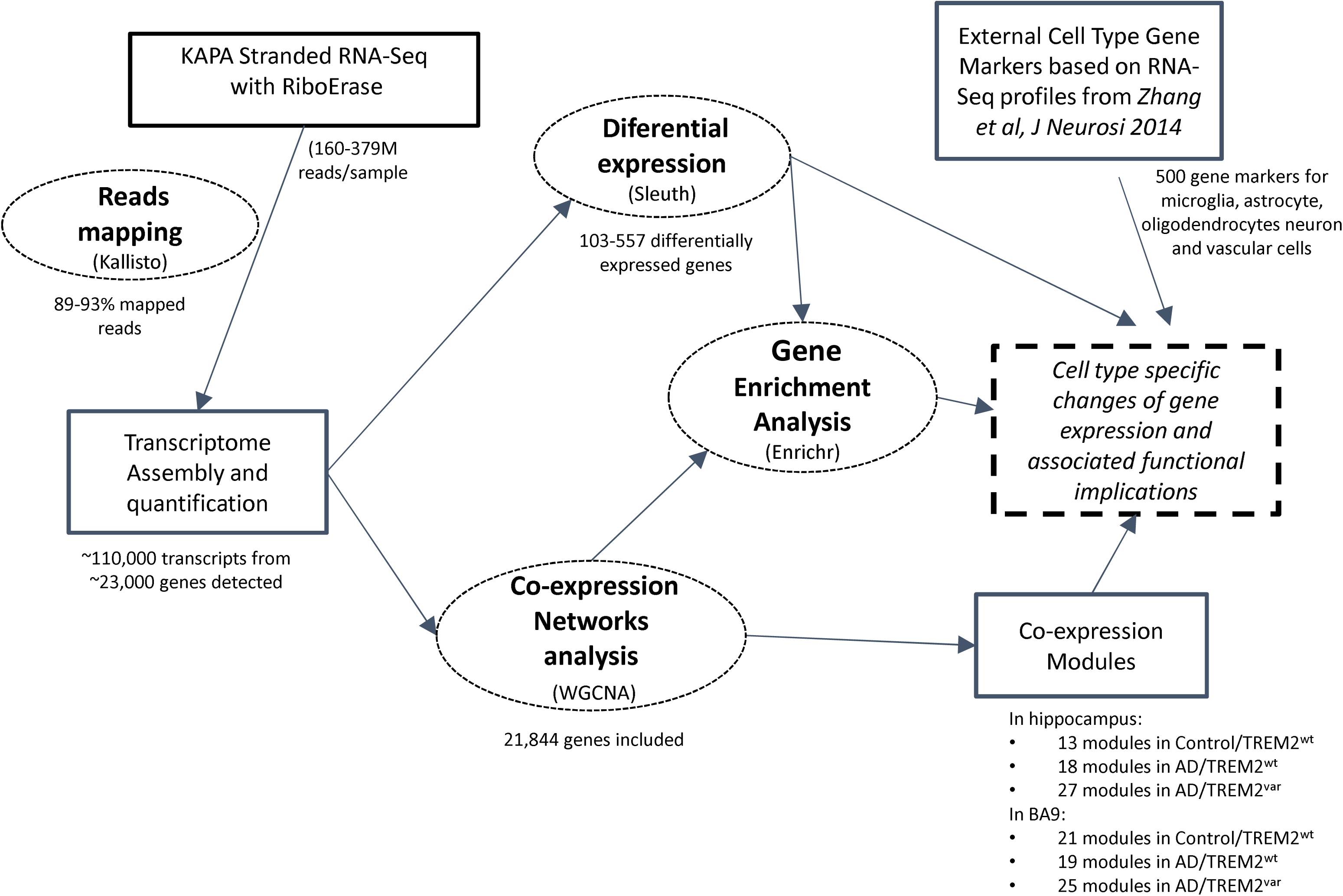
Data analysis pipeline. Transcriptome profiling of 69 brain donors (131 samples) comparing **Control_MCI_AD/TREM2^var^**, **AD/TREM2^wt^** and **Control/TREM2^wt^** from hippocampus and BA9 tissue using RNA-Seq. Read alignment to quantify transcripts was achieved with Kallisto and different expression determined using Sleuth. Coexpression network analysis was carried out using WGCNA. Enrichment analysis used Enrichr. To estimate cell type associations with coexpression modules, gene markers were obtained from an external source (Zhang *et al*., 2014). Results were contrasted to infer AD and tissue-specific changes caused by the presence of a disease-associated TREM2 variant.

### TREM2 cohort and RNA extraction

Informed consent for all brain donors was obtained according to the Declaration of Helsinki (1991) and protocols and procedures were approved by the relevant ethical committee and obtained by each brain bank. Pathological diagnosis was made according to established methods at the time of donation (Braak and Braak, 1991; Braak *et al*., 2003; Alafuzoff *et al*., 2006; Alafuzoff *et al*., 2008; Josephs *et al*., 2014). Where necessary for historical cases, retrospective diagnoses were made using current criteria. Tissue from a total of 69 brain donors (131 samples) were obtained for the following three groups: **Control/MCI/AD/TREM2^var^** (donors with a non-synonymous DNA variant in TREM2 expected to put them at higher risk of AD), **AD/TREM2^wt^** (AD donors with no TREM2 disease associated variant) and **Control/TREM2^wt^** (tissue from age-matched donors with no disease associated TREM2 variant nor any evidence of neurological problems or AD pathology) (Table 1 and Table S1).

**Table 1:**
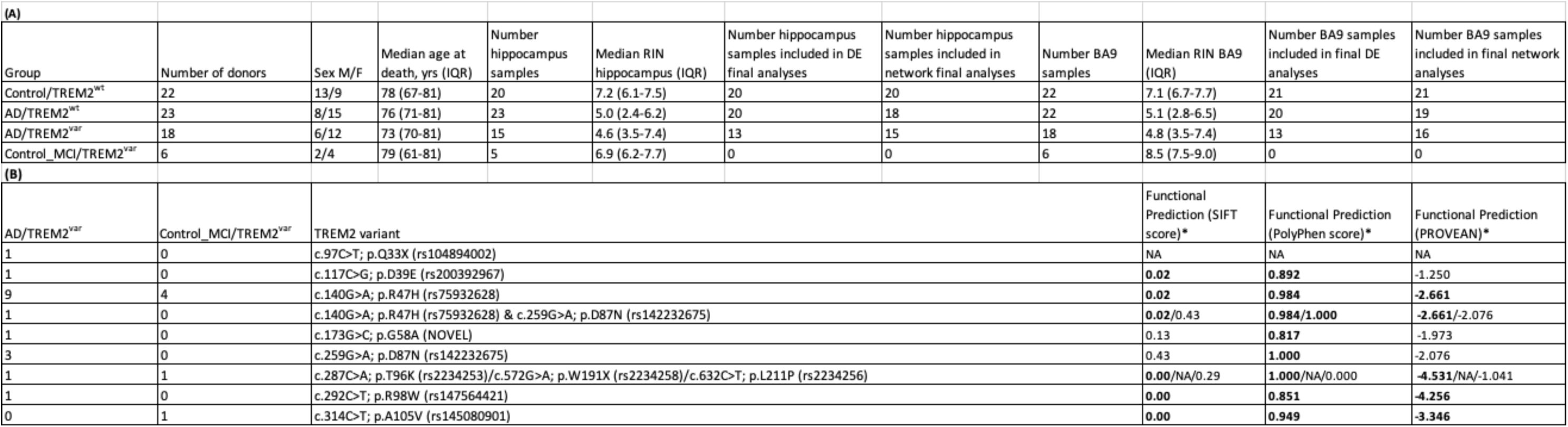
Cohort Characteristics. **(A)** Number, sex and age of donor hippocampus and BA9 tissue used in the study. A total of 69 brain donors (131 samples) were obtained for the following three groups; **Control_MCI_AD/TREM2^var^** (donors with a non-synonymous DNA variant in TREM2 expected to put them at higher risk of AD), **AD/TREM2^wt^** (AD donors with no TREM2 disease associated variant) and **Control/TREM2^wt^** (tissue from age-matched donors with no disease associated TREM2 variant nor any evidence of neurological problems or AD pathology). **(B)** The majority of **TREM2^var^ cases** have c.140G>A; p.R47H (rs75932628) variant or other variant previously described in AD patients. c.173G>C; p.G58A is a novel variant. *All TREM2 variants are predicted to have functional consequences by either SIFT, PolyPhen and/or PROVEAN) based on TREM2 coding transcript ENST00000373113.7 (Q5TCX1 Q9NZC2).

The Control/MCI/AD/TREM2^var^ group consisted of 24 donors: 10 donors were identified by sequencing TREM2 Exon 2 in 631 AD and normal elderly control brain donors from the MRC London Neurodegenerative Diseases Brain Bank (King’s College London, UK); 4 donors were identified from a screen of 198 AD cases from The Netherlands Brain Bank (Royal Netherlands Academy of Arts and Sciences, Netherlands) and a further 10 donors which have been described previously (Guerreiro *et al*., 2013a) were obtained from the Thomas Willis Brain Collection (John Radcliffe Hospital, UK) (4 donors) and from the Queen Square Brain Bank for Neurological Disorders (University College London, UK) (6 donors). Within the Control/MCI/AD/TREM2^var^ group, 18 had a pathologically confirmed diagnosis of AD while 6 were pathologically normal at death, although one donor had symptoms of mild cognitive impairment just prior to death and their post mortem pathology was assessed as Braak stage III (Table S1). As expected, the majority of Control/MCI/AD/TREM2^var^ cases had the variant c.140G>A; p.R47H (rs75932628), while the remainder had one of eight additional variants in TREM2 predicted to be detrimental. One donor had a novel variant while the remainder have been described previously in AD cases (Table 1 and Table S1).

Where available, ∼100 mg of frozen tissue from post mortem hippocampus and pre-association cortex (BA9) from each donor, reflecting early and later affected areas in AD pathogensis (Mastrangelo and Bowers, 2008; Matarin *et al*., 2015), were homogenized in matrix-lysing D tubes containing Qiazol reagent. RNA was isolated using the miRNeasy Mini Kit (Qiagen). Both total RNA and small RNA fractions were extracted separately according to the manufacturer’s instructions. The quantity and quality of total RNA was assessed using the NanoDrop 8000 spectrophotometer V2.0 (ThermoScientific, USA) and Agilent 2100 Bioanalyser (Agilent Technologies, Waldbronn, Germany), respectively. As reported previously for post mortem brain samples (Proitsi *et al*., 2014), RNA integrity numbers (RIN) were variable, ranging from 1.2 to 9.5 with a median of 6.3 (Table 1 and Table S1).

### Library generation and RNA-sequencing

100ng of total RNA from each sample was used to prepare total RNA libraries using the KAPA Stranded RNA-Seq Kit with RiboErase (KAPA Biosystems, Massachusetts, USA). Fragmentation prior to first strand cDNA synthesis was carried out using incubation conditions recommended by the manufacturer. For degraded samples (those with a RIN<7) 65°C for 1 minute was used, while 94°C for 6 minutes was used for the remaining samples. 14 cycles of PCR were performed for final library amplification. Resulting libraries were quantified using the Qubit 2.0 spectrophotometer (Life Technologies, California, USA) and average fragment size assessed using the Agilent 2200 Tapestation (Agilent Technologies, Waldbronn, Germany). Equimolar amounts of each library with compatible indexes were pooled together for sequencing, resulting in 5 pools with 22 samples, and one pool of 23. 75bp paired-end reads were generated for each library pool using the Illumina NextSeq®500 (Illumina Inc., Cambridge, UK) in conjunction with the NextSeq®500/550 High-Output 150cycle kit. Each pool was sequenced a total of 6 times in order to achieve 100 million reads per sample. Expression data generated in this study are available at NCBI’s GEO through accession number GSEXXXXXX.

### RNA-Seq Quality Control and Differential Expression

RNAseq reads were aligned to the human GRCm38 rel 79 reference transcriptome using Kallisto version 0.42.4 (Bray *et al*., 2016). This transcriptomes was constructed using Ensembl reference transcriptomes (https://www.ensembl.org/info/data/ftp/index.html.) We downloaded the reference transcriptome from the Kallisto project website (“http://bio.math.berkeley.edu/kallisto/transcriptomes/”). The aforementioned resource provided by the Kallisto project is not available anymore and reference transcriptomes can now be downloaded from the patcherlab github at https://github.com/pachterlab/kallisto-transcriptome-indices/releases where further detail can be obtained on how the indexes are generated. The efficiency of the mapping varied from 79 to 94% with the number of reads mapped ranging from 159 million to 379 million (Table S1).

We applied a number of quality control methods implemented in the R Sleuth package. Principal component analysis revealed that samples clustered by RIN score, so we included RIN score as a covariate in Differential Expression (DE) analysis or removed samples with a RIN score <2.2 for network analysis, where this variable could not be controlled (Table S1 and Supplementary File 1). We aligned the RNAseq reads using STAR version 2.5.2a and called variants for the Y chromosome using samtools version 1.3+htslib-1.3 and bcftools 1.3. Reassuringly, the number of variants called were always higher in the males when we checked against their recorded sex, except for one sample which had also failed to meet sequencing quality and was therefore excluded from the analysis. We also confirmed the presence of the expected TREM2 variant in the relevant reads as well as conordance of their APOE genotype determined independently. Overall, there was high sample concordance and high sequencing quality. Only a small number of samples were excluded from further analysis due to low number of reads count (implying sequencing failure) and/or because they were outlier samples relative to the population norm and/or because they had a high rate of duplicate reads and/or where samples with the TREM2 variant did not have a diagnosis of AD (Table S1). Overall, this left 53 hippocampus and 54 BA9 samples for DE analysis and 53 hippocampus and 56 BA9 samples for network analysis (Table 1).

Following quality control filtering, in hippocampus 109,968 transcripts representing 24,467 genes were available for the **Control/TREM2^wt^** and **AD/TREM2^wt^** comparison and 109,569 transcripts representing 24,403 genes were available for the **Control/TREM2^wt^** and **AD/TREM2^var^** comparison. In BA9, 111,281 transcripts representing 24,782 genes were available for the **Control/TREM2^wt^** and **AD/TREM2^wt^** comparison and 110,669 transcripts representing 24,717 genes were available for the **Control/TREM2^wt^** and **AD/TREM2^var^** comparison. We performed DE analysis on TPM normalized counts with the R sleuth package version 0.28.1 (Pimentel *et al*., 2017). We used Wald tests provided by the sleuth package to statistically compare expression differences in **Control/TREM2^wt^** and **AD/TREM2^wt^** and **Control/TREM2^wt^** and **AD/TREM2^var^**, controlling for differences in RIN score, sex and age at death by including their values as covariates in the sleuth model.

We generated plots and graphics using the sleuth (H. Pimentel et al., 2016; H. J. Pimentel et al., 2016) and ggplot version 2.2.1 (Wickham, 2009) R packages. We used R version 3.3.0.

### Comparison in differential gene expression between Control/TREM2^wt^ and AD/TREM2^var^ versus Control/TREM2^wt^ and AD/TREM2^wt^

To assess if differences in gene expression between **Control/TREM2^wt^** and **AD/TREM2^var^** are recapitulated between **Control/TREM2^wt^** and **AD/TREM2^wt^**, we assessed the degree of overlap in the lists of differentially expressed genes for each comparison in hippocampus and BA9 using a similar approach to (Kuhn *et al*., 2007).

We selected the top ranked isoform of each gene by p-value (q-value <0.05) from the **Control/TREM2^wt^** and **AD/TREM2^wt^** analysis and compared its ranking in the **Control/TREM2^wt^** and **AD/TREM2^var^** comparison. Genes were ‘condordant’ if they changed in the same direction in both comparisons (same b value sign). There was strong over-representation of concordant changes in the top ranking genes across in both comparisons (Supplementary File 1).

To establish the degree of the association between the two lists of differentially expressed genes we defined a ‘concordance coefficient’. The concordance coefficient was calculated as:

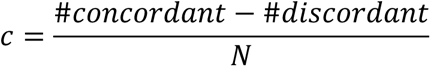

where #concordant is the number of concordant genes between the two comparisons, #discordant is the number of discordant genes and N the number of genes selected for the comparison. The value of c ranges from 1 for a perfectly concordant set to -1 for a completely discordant one. A value of 0 reflects the same number of concordant and discordant genes. We obtained a value of 1 for both Entorhinal Cortex and BA9 comparisons (i.e. perfect concordance).

To determine the strength of the association we used three approaches:

1. We permutated the labels of the samples for **Control/TREM2^wt^** and **AD/TREM2^var^** 100 times and compared this to the original outcome. Not one of the alternative comparisons was as strongly associated as the original (i.e. pval < 0.01 for the strength of the association)
2. To test the specificity of the association to the AD signature we repeated the permutation for **AD/TREM2^wt^** vs **Control/TREM2^wt^** and compared this against the original **Control/TREM2^wt^** and **AD/TREM2^var^** signature. The result was the same as before (i.e. pval < 0.01 for the strength of the association)
3. Finally, to further test how strongly the **Control/TREM2^wt^** and **AD/TREM2^var^** signature recapitulates the **AD/TREM2^wt^** vs **Control/TREM2^wt^** result, we compared it to a random **AD/TREM2^var^** vs **Control/TREM2^wt^** gene list. Random signatures were generated by drawing (without replacement) groups of N non-redundant genes from genes matched between the two comparisons. We generated a concordant coefficient for each of 10000 random **AD/TREM2^var^** vs **Control/TREM2^wt^** signatures. In all cases the permutated coefficient was significantly smaller than the concordance coefficient observed between **AD/TREM2^var^** vs **Control/TREM2^wt^** and **AD/TREM2^wt^** vs **Control/TREM2^wt^**.

### Gene ontology and pathway enrichment analysis

Genes with transcripts reaching significance, q-value ≤ 0.01, were used for gene ontology and pathway analysis. We performed gene ontology and pathway analysis using the Enrichr database (Chen *et al*., 2013; Kuleshov *et al*., 2016) as described previously (Newhouse, 2017).

### Gene network construction and module detection

For the co-expression analysis, we selected the transcript isoform with highest mean expression in the filtered list to represent each gene. Genes where there were at least 10 reads in more than 90% of samples for each tissue or group were included (Langfelder and Horvath, 2008). This left us with 21,844 genes (Figure 1).

We used WGCNA version 1.51 to identify modules of co-expressed genes within gene expression networks (Langfelder and Horvath, 2008). To construct the networks, we calculated biweight midcorrelations for all possible genes pairs. We entered the values in a matrix, and transformed the data so that the matrix followed an approximate scale-free topology (Supplementary File 1). We used a dynamic tree cut algorithm to detect network modules and used signed networks to preserve the direction of correlations. We ran singular value decomposition on each module’s expression matrix and used the resulting module eigengene, which is equivalent to the first principal component (Langfelder and Horvath, 2007), to represent the overall expression profiles of the modules.

We used a hypergeometric test to match modules with the most significant gene overlap between **Control/TREM2^wt^** and **AD/TREM2^wt^**, and, **Control/TREM2^wt^** and **AD/TREM2^var^** for each brain region. The **AD/TREM2^wt^** and **AD/TREM2^var^** groups were subsequently re-coloured to match the corresponding colour used for each module in the **Control/TREM2^wt^** group. Modules with no significant overlap retained their original colour since these modules don’t appear in the comparator group.

### Module preservation statistics

We applied both the module preservation Zsummary and the medianRank statistics to assess the module preservation between groups (Langfelder *et al*., 2011). Unlike the cross-tabulation test, Zsummary not only considers the overlap in module membership, but also the density and connectivity patterns of modules. We converted the transcript-level measurements into gene-level measurements using the collapseRows R function and adapted the method to select the most representative transcript by choosing the transcript isoform with the highest average expression to represent that gene, as this has been shown to have the highest between-study consistency (Miller *et al*., 2011). Overall, we retained 21,844 genes for further analysis. We then used student t-tests to statistically compare expression differences between modules as suggested by the WGCNA authors (Langfelder and Horvath, 2008).

### Module membership statistics

The level of correlation between each gene and each module eigenene was assessed to determine how strong the connection of a gene was with the rest of the module it had been assigned to, and to each of the other modules. Two different approaches were used, based on the methods described by the WGCNA authors (Langfelder and Horvath, 2008). The first approach used all genes to determine overall conservation of the modules. The second used only those genes assigned to a given module to assess “hub” conservation i.e. this assesses if genes more strongly correlated to a given module in one of the networks (e.g. Control/TREM2^wt^) are strongly correlated to the equivalent module in a comparator network (e.g. AD/TREM2^wt^).

We used Cytoscape v3.7.0 to generate network maps of the most highly correlated genes and modules for each group, retaining the same colours used for each module generated in WGCNA (Shannon *et al*., 2003). We only display the genes with correlations stronger than 0.5 for clarity. This enables graphic visualisation of relationships between modules and how they change between the three groups and brain regions being compared (Control/TREM2^wt^, AD/TREM2^wt^ and AD/TREM2^var^).

### Integration of cell type markers

A recent study showed that there is a large degree of conservation in cell type-specific transcriptome-wide RNA expression in brain tissue (McKenzie *et al*., 2018). This conservation is not only high between independent studies, but is also high between equivalent cells in human and mouse brain, thus supporting the validity of using the cell type dataset for the current study. We therefore selected the top 500 most highly expressed genes for different brain cell types (astrocytes, microglia, neurons, endothelial cells, oligodendrocyte precursor cells, newly formed oligodendrocytes and myelinating oligodendrocytes) from a mouse RNA-Sequencing transcriptome database (Zhang *et al*., 2014). We obtained human orthologs of these genes using biomart (http://www.biomart.org/). Of these, it was possible to map 419 marker genes from astrocytes, 456 from endothelial cells, 447 from microglia, 431 from myelinating oligodendrocytes, 431 from newly formed oligodendrocytes, 420 from oligodendrocyte precursors cells and 439 from neurons, to our dataset. These were used in subsequent analyses. To compare cell type enrichment for each module, the total number of markers present in a given module was divided by the total number of markers available for each cell type to normalise and enable comparison between different sized modules. These values were then multipled by 100 to avoid working with very small numbers.

## Results

### Differentially expressed transcripts in AD largely AD rather than TREM2-dependent

First, we explored the number and directionality of differentially expressed transcripts (DE) for hippocampus and BA9 between AD cases (**AD/TREM2^wt^**) and AD cases with a disease-associated TREM2 variant (**AD/TREM2^var^**) compared to age-matched controls (**Control/TREM2^wt^**) (Tables S2-S5). Around 110,000 transcripts representing ∼23,000 genes were included in this analysis. As expected, there were more DE transcripts in hippocampus compared to BA9 in AD (qval<0.05) and greater numbers of up- than down-regulated DE transcripts in AD cases in both tissues (Figure 2). However in AD with a TREM2 variant, there were not only fewer DE genes overall in both tissues compared to AD (although as with non TREM2 carrier AD, BA9 was less affected) but greater numbers of down-regulated rather than upregulated DE genes reaching nominal significance (qval<0.05) (Figure 2).

**Figure 2:**
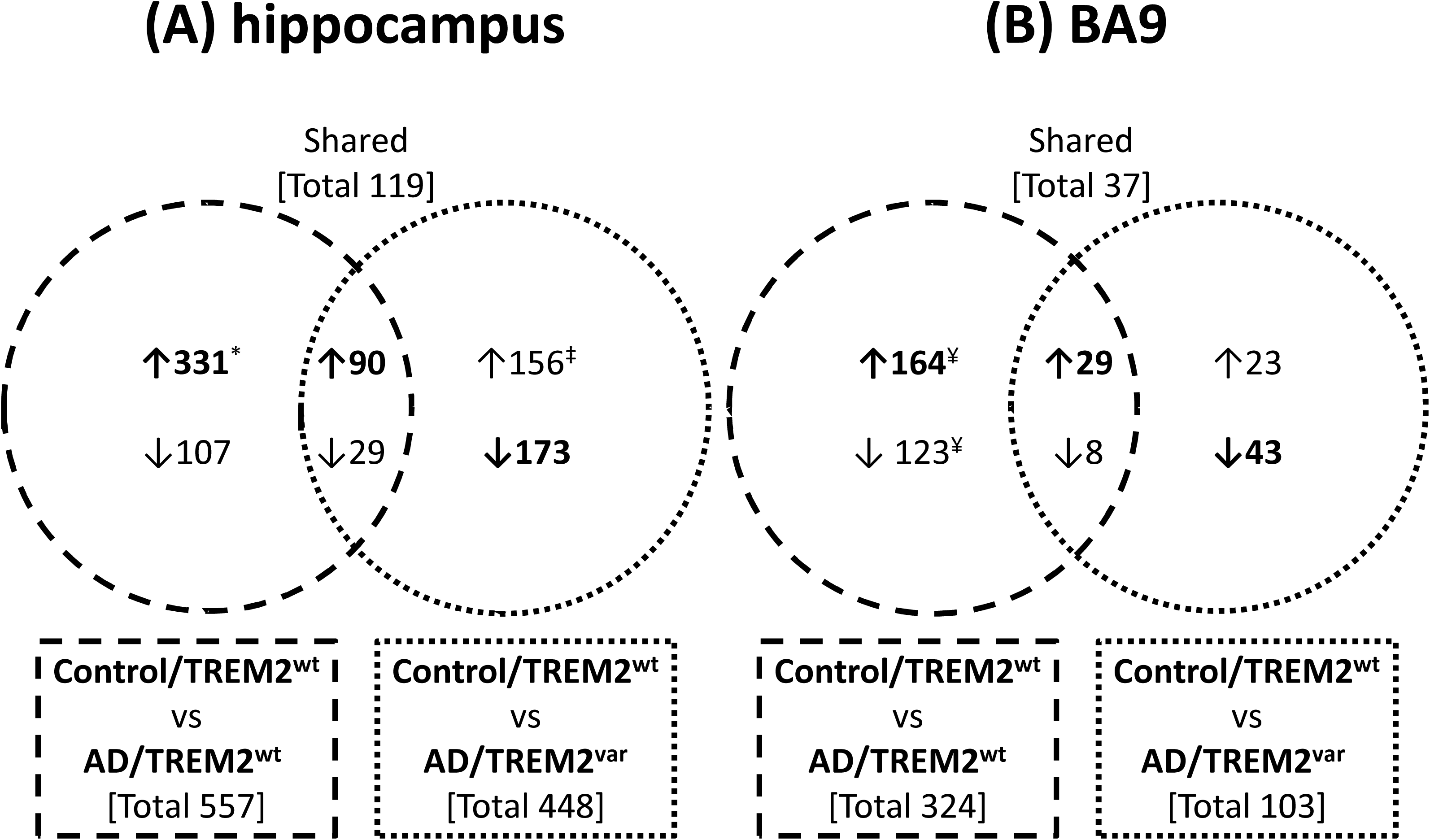
The number of differentially expressed transcripts (qval<0.05) in **Control/TREM2^wt^ vs AD/TREM2^wt^** and **Control/TREM2^wt^ vs AD/TREM2^var^** in hippocampus **(A)** and BA9 **(B)** including those upregulated (↑) and downregulated (↓) and overlapping between both AD groups. 108,298-110,012 transcripts were detectable in hippocampus and BA9 across the three diagnostic groups. Of those meeting qval<0.05, 8 transcripts were below detection in AD/TREM2^var^ (*) and 2 transcripts were below detection in AD/TREM2^wt^ (‡) in hippocampus and 2 transcripts were below detection in AD/TREM2^var^ (¥) in BA9.

This likely reflects differences in the number of samples available for each group for analysis, as although the numbers of transcripts reaching nominal significance for DE (qval <0.05) was generally lower in those with a TREM2 variant, there was remarkable concordance in the identity and direction of change of the top ranking differentially expressed transcripts and genes, in both tissue area’s and AD groups. This is unsurprising given both groups of AD have common pathological hallmarks and indeed, are indistiguishable at the microscopic level. The concordance coefficient in DE was 1 in both tissues, indicating a high degree of similarity. In further tests comparing concordance against randomly selected DE genes, the permutated coefficients were always lower than the actual concordance coefficient for both tissues.

However, embedded amongst the many common molecular changes present in end stage AD, we were particularly interested in uncovering TREM2-dependent differences which might signify cells, processes, pathways and genes directly controlled by TREM2 and therefore impacted by TREM2 dysfunction, which could be important targets for treatment development. TREM2- dependent genes were expected to be those altered between **Control/TREM2^wt^** and **AD/TREM2^wt^**, but not different between **Control/TREM2^wt^** and **AD/TREM2^var^** or having an exacerbated response in **AD/TREM2^var^**. Examples of such DE genes in hippocampus were LILRB4 and LSR which were significantly up-regulated only in AD cases without a TREM2 variant (Figure 3A and B). Many transcripts displaying this pattern had very low levels of expression, and sometimes fell below the level of detection in TREM2 carriers, thus explaining their absence from the DE list in **Control/TREM2^wt^** and **AD/TREM2^var^** (Tables S2 and 3). Alternatively, genes such as LMX1B and PUF60 were down-regulated in AD, but only in donors without a TREM2 variant (Figure 3C and D, Tables S2 and 3) while SORBS1 and SNX22 had a noticeably greater magnitude of DE in AD cases with a TREM2 variant (Figure 3D and E, Tables S2 and 3). Many of the TREM2- dependent DE genes appear to have enriched expression in oligodendrocytes in the human brain (McKenzie *et al*., 2018) and/or are ascribed roles in microtubule and cytoskeletal activity.

**Figure 3:**
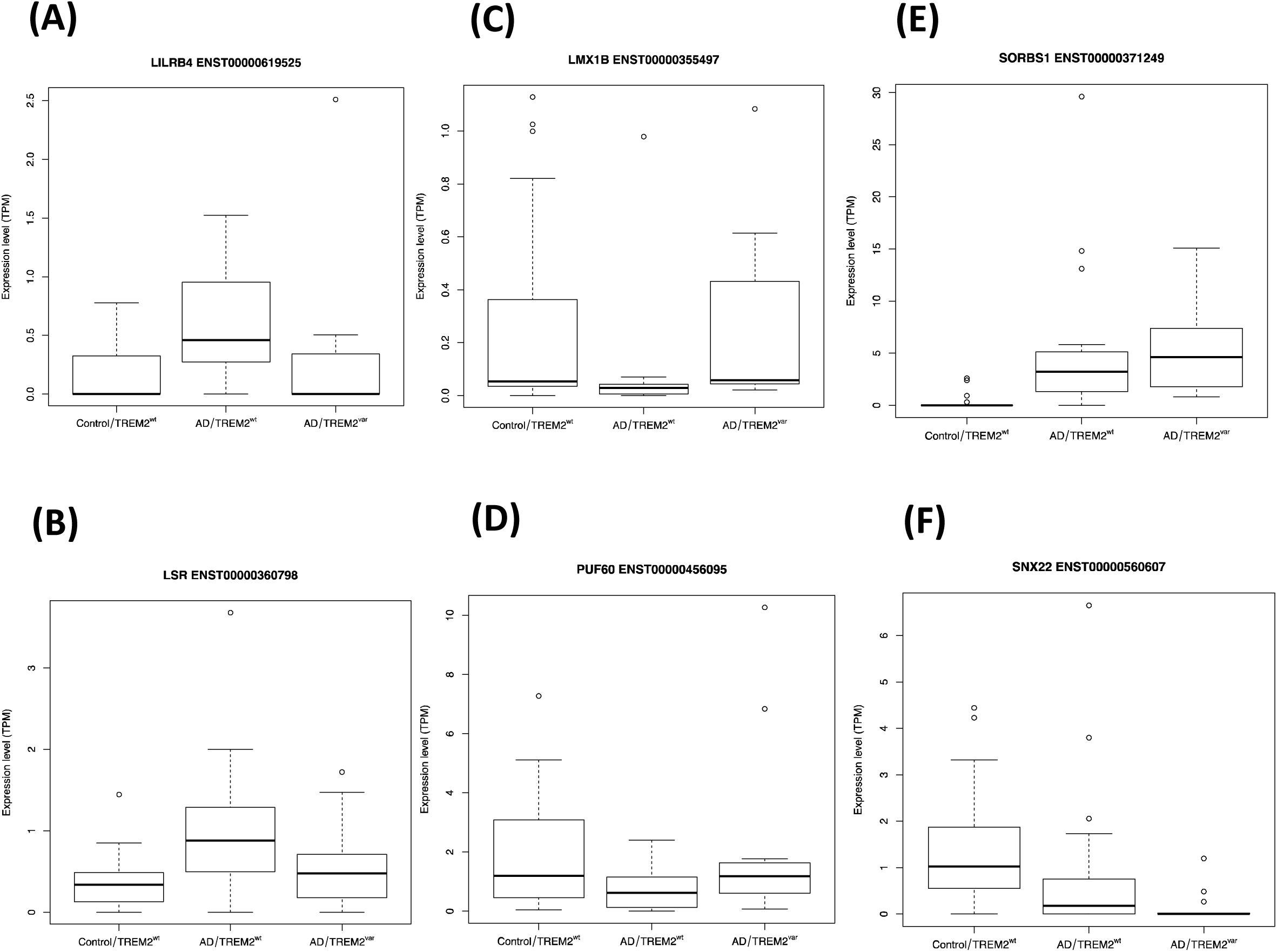
Examples of TREM2-dependent genes (transcripts) displaying differential expression between **Control/TREM2^wt^** and **AD/TREM2^wt^**, but either not displaying differential expression between **Control/TREM2^wt^** and **AD/TREM2^var^** or having an exacerbated response in **AD/TREM2^var^**. LILRB4 **(A)** and LSR **(B)** were significantly up-regulated in hippocampus only in AD cases without a TREM2 variant. LMX1B **(C)** and PUF60 **(D)** were significantly down-regulated in hippocampus only in AD cases without a TREM2 variant. SORBS1 (E) and SNX22 (F) had noticeably greater or less differential expression, respectively, in AD cases with a TREM2 variant

### TREM2 and TYROBP central to co-expression module in AD hippocampus

To better understand the impact of TREM2 dysfunction within or between closely coordinated groups of genes within the brain, we next generated hippocampus and BA9 co-expression networks for each group (**Control/TREM2^w^**^t^, **AD/TREM2^wt^** and **AD/TREM2^var^**) (see Methods and Figure 1). The hippocampus **Control/TREM2^wt^** co-expression network consisted of 14 modules, each containing between 7 and 9770 gene members, while the **AD/TREM2^wt^** and **AD/TREM2^var^** modules consisted of 19 modules (6 to 11957 genes) and 28 modules (1 to 10768 genes), respectively (Figure 4). The bisque4 module contains the largest number of genes, while the grey module contains unassigned genes and was therefore excluded from further consideration. Similarly, the **BA9 Control/TREM2^wt^** co-expression network consisted of 22 modules, each containing between 15 and 8126 gene members, while the **AD/TREM2^wt^** and **AD/TREM2^var^** modules consisted of 20 modules (10 to 11677 genes) and 26 modules (2 to 6837 genes), respectively (Figure S1).

**Figure 4:**
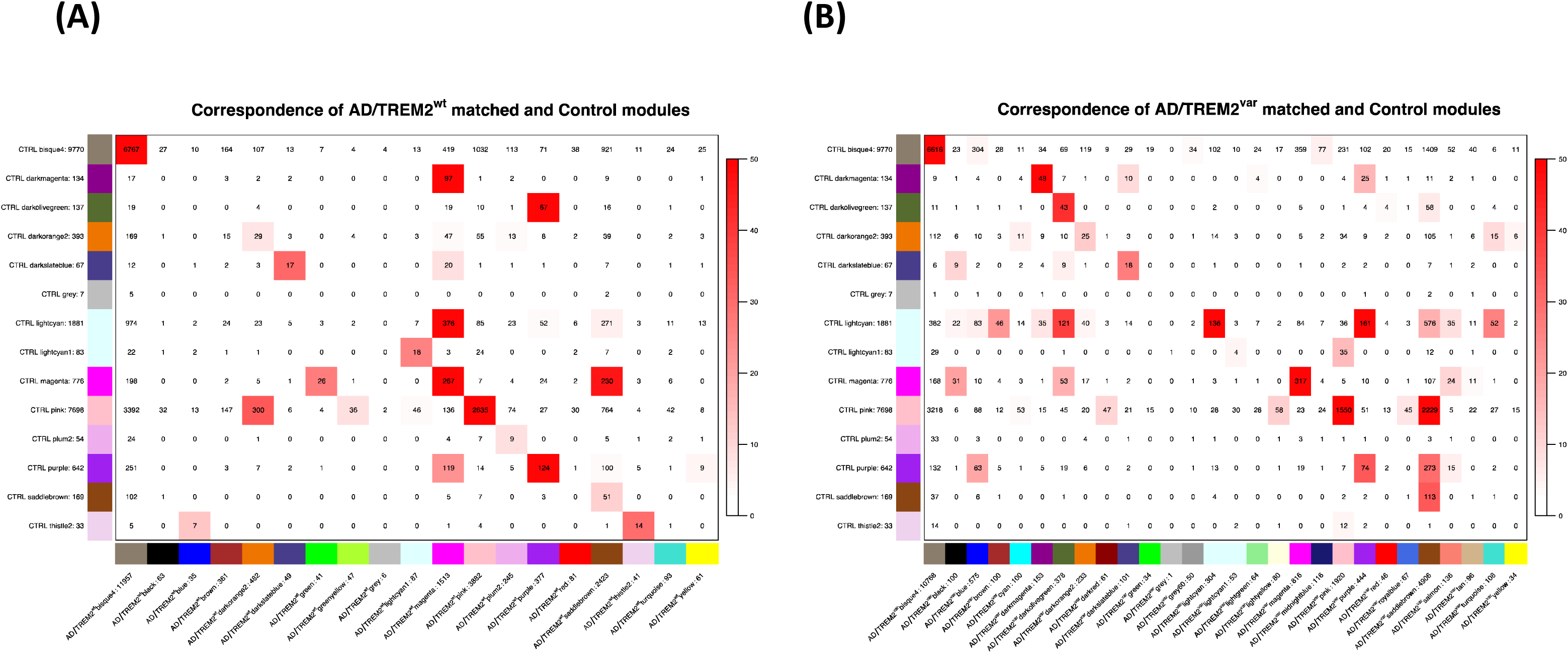
WGCNA modules and their preservation between **Control/TREM2^wt^** and **AD/TREM2^wt^** networks **(A)** and **Control/TREM2^wt^** and **AD/TREM2^var^** networks **(B)** in hippocampus. Each row of the table corresponds to one **Control/TREM2^wt^** set-specific module (assigned a colour as indicated by the text label). Each column corresponds to one **AD/TREM2^wt^** or **AD/TREM2^var^**set-specific module, respectively. Numbers indicate the number of genes in each module which are also divided across the table to indicate the number of genes which intersect each pair of modules between the two groups. Red shading within the table is generated by $-*∖*log(p)$, with p being the hypergeometric test p-value for the degree of overlap of each module pair. The stronger the red color, the more significant the overlap is. Most **Control/TREM2^wt^** set-specific modules have a corresponding **AD/TREM2^wt^** counterpart or involve a split or merger involving only two modules **(A)**. In contrast, most of the **Control/TREM2^wt^** set-specific modules are not conserved in **AD/TREM2^var^**, where there is significant fragmentation and generation of additional modules **(B)**. The grey module which contains unassigned genes is excluded from consideration.

In hippocampus, most **Control/TREM2^w^**^t^ modules had an **AD/TREM2^wt^** counterpart, or involved the split or merger of genes across only two modules. The exception was the **Control/TREM2^w^**^t^ dark magenta module, which expanded in **AD/TREM2^wt^** to take in genes from 5 different **Control/TREM2^w^**^t^ modules (including magenta and dark magenta) to create a new magenta module, and in so doing became the third largest module in **AD/TREM2^wt^** (Figure 4A). Two other notable changes included around a third of the magenta module genes in **Control/TREM2^w^**^t^ splitting to join the saddle brown module in **AD/TREM2^wt^** and half of the dark olive green module genes from **Control/TREM2^w^**^t^ joining the purple module in **AD/TREM2^wt^**, thus causing the dark olive green module to disappear as an independent module in **AD/TREM2^wt^**. Interestingly, the purple module contained TREM2 and TYROBP in both **Control/TREM2^w^**^t^ and **AD/TREM2^wt^** hippocampus (Table S6 and S7). In contrast, there was greater fragmentation in **AD/TREM2^var^** where very few modules in **Control/TREM2^w^**^t^ were preserved in **AD/TREM2^var^**. Over half of all **Control/TREM2^w^**^t^ modules became split between more than two **AD/TREM2^var^** modules (Figure 4B). Unlike what was found in **AD/TREM2^wt^** (Figure 4A) the magenta module remained largely preserved in **Control/TREM2^w^**^t^ and **AD/TREM2^var^** (Figure 4B). The purple module in **Control/TREM2^w^**^t^ on the other hand, instead of taking in dark olive green genes as was found in **AD/TREM2^wt^**, fragmented to join blue, saddlebrown and purple modules in **AD/TREM2^var^** (Table S8).

### TREM2 and TYROBP lose their module influence in AD hippocampus in TREM2 variant carriers

TREM2 and TYROBP completely lost their strong position as intermolecular hub genes with high module membership (MM) ranking in AD TREM2 carriers compared to the very central position they had in the purple module in control and AD hippocampus. While TREM2 had the 12^th^ highest module membership rank in the purple module in control hippocampus (MM correlation 0.92) and 29^th^ in AD (MM correlation 0.93), TYROBP was 48^th^ (MM correlation 0.88) and 4^th^ (0.96), respectively (Table S6 and S7). Module membership ranking dropped to a lowly 6,983^rd^ (MM correlation 0.62) for TREM2 and 9,861^st^ position for TYROBP (MM correlation 0.51) in the saddlebrown module they were assigned to in AD TREM2 carriers.

Some modules were better preserved than others, indicating member genes had retained a tightly coordinated pattern of gene expression. Of note, the magenta module in the hippocampus in **Control/TREM2^w^**^t^ had the lowest median rank score amongst the modules and was one of three modules with a z score >10 in **AD/TREM2^wt^** and **AD/TREM2^var^**, indicating high module preservation (Figure S2A-D). Although less strong, a similar profile for the pink and purple modules was also found in hippocampus. In BA9, high module preservation was consistently found for the darkgreen and lightpink4 modules (Figure S2E-H).

In BA9, the majority of **Control/TREM2^w^**^t^ modules had an **AD/TREM2^wt^** counterpart or consisted of genes splitting between only two modules in **AD/TREM2^wt^**, suggesting overall a relatively mild impact of AD at the level of gene expression in this tissue (Figure S1A). There were a few notable exceptions where related modules behaved differently in the **AD/TREM2^var^** group, suggesting some AD impacted groups of genes were dependent on TREM2 (Figure S1B). The black module in **AD/TREM2^wt^** was largely formed by the merger of genes from the black and dark green modules in **Control/TREM2^w^**^t^ while in **AD/TREM2^var^**, rather than joining the black module, many of the dark green module genes instead moved to a new salmon module not present in **AD/TREM2^wt^**. The greenyellow module was the third largest module in **Control/TREM2^w^**^t^ and contained TREM2 and TYROBP (Table S9). In **AD/TREM2^wt^**, this module fragmented in to blue (containing TREM2 and TYROBP, Table S10), dark red and green yellow modules while in **AD/TREM2^var^**, it fragmented in to blue, dark red and greenyellow modules (containing TREM2 and TYROBP, Table S11) as well as a new sky blue 2 module which also drew genes from the steelblue and green modules found in **Control/TREM2^w^**^t^. In contrast, the steelblue and green modules remained largely unchanged in **Control/TREM2^w^**^t^ and **AD/TREM2^var^** suggesting AD pathology is unable to impact these genes in the absence of fully functional TREM2. Interestingly, TREM2 and TYROBP had very little influence in the greenyellow module in control where AD pathology is absent or in AD TREM2 carriers as would be expected from TREM2 dysfunction. In contrast, TREM2 and TYROBP had significant influence in the blue module in AD, where they were ranked 19^th^ (MM correlation 0.90) for TREM2 and 40^th^ (0.85) for TYROBP (Tables S9, S10 and S11).

### TREM2 and TYROBP in hippocampus module with microglia identity with influence of oligodendrocyte and endothelial modules in AD/TREM2^wt^ but not in AD/TREM2^var^

We next investigated whether any of the modules had a particular cell type identity by looking for over-representation of genes previously identified as having highly enriched or exclusive expression in specific brain cell types (endothelial cells, microglia, neurons, astrocytes, newly formed oligodendrocytes, oligodendrocyte precursor cells, myelinating oligodendrocytes).

As noted above, the purple module in hippocampus contained TREM2 and TYROBP in both **Control/TREM2^w^**^t^ and **AD/TREM2^wt^** groups and along with other genes gave this module a very clear microglial identity in control and AD (Figure 5A). No other module in either group had such a distinctive microglia identity. However, this module became fragmented in **AD/TREM2^var^** to form the blue and saddlebrown modules. Although both retained a weaker microglia identity and TREM2 and TYROBP still resided in the new saddlebrown microglia module (Figure 5A), they both dramatically lost their central influence over this module (Figure 5A).

**Figure 5:**
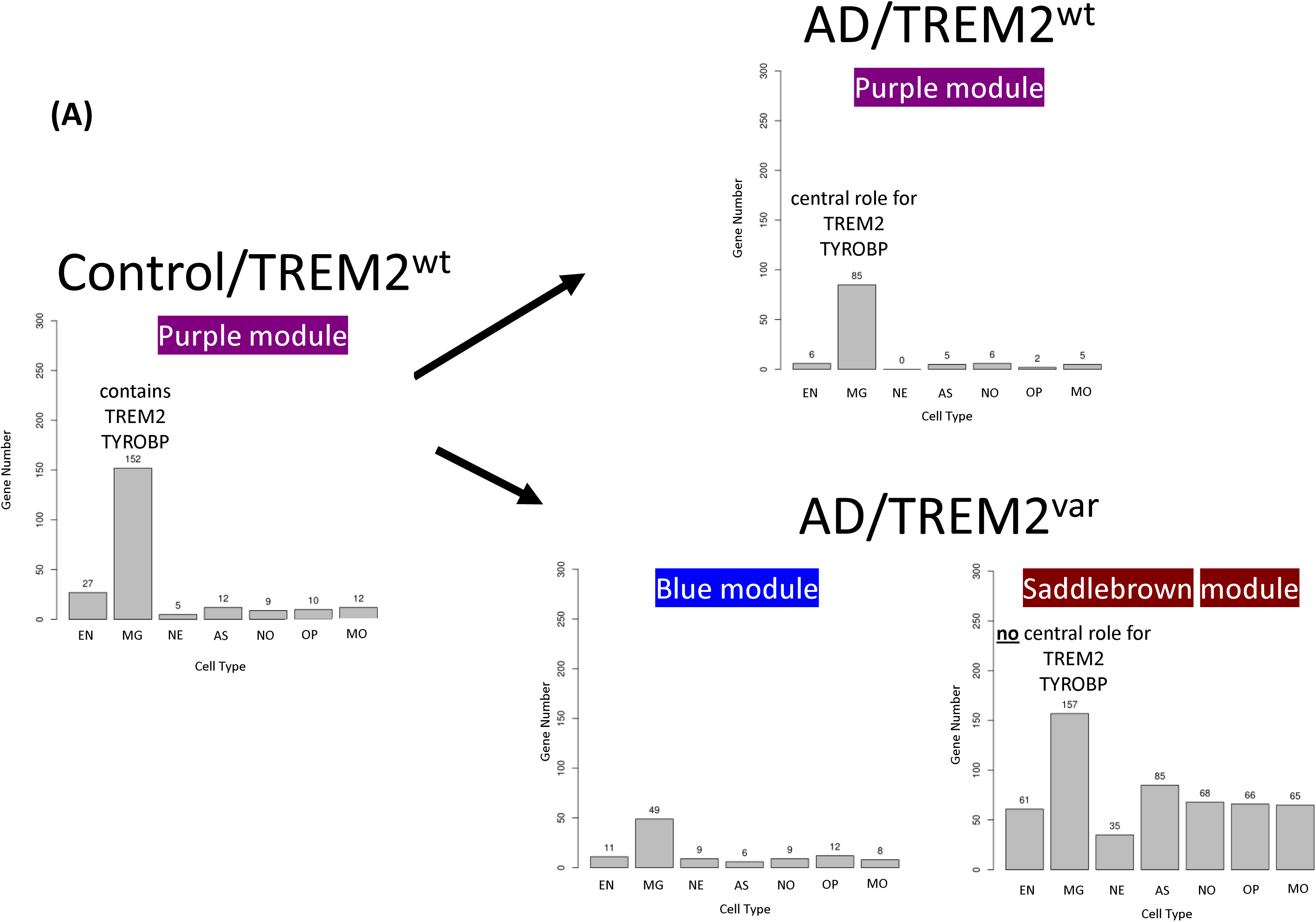
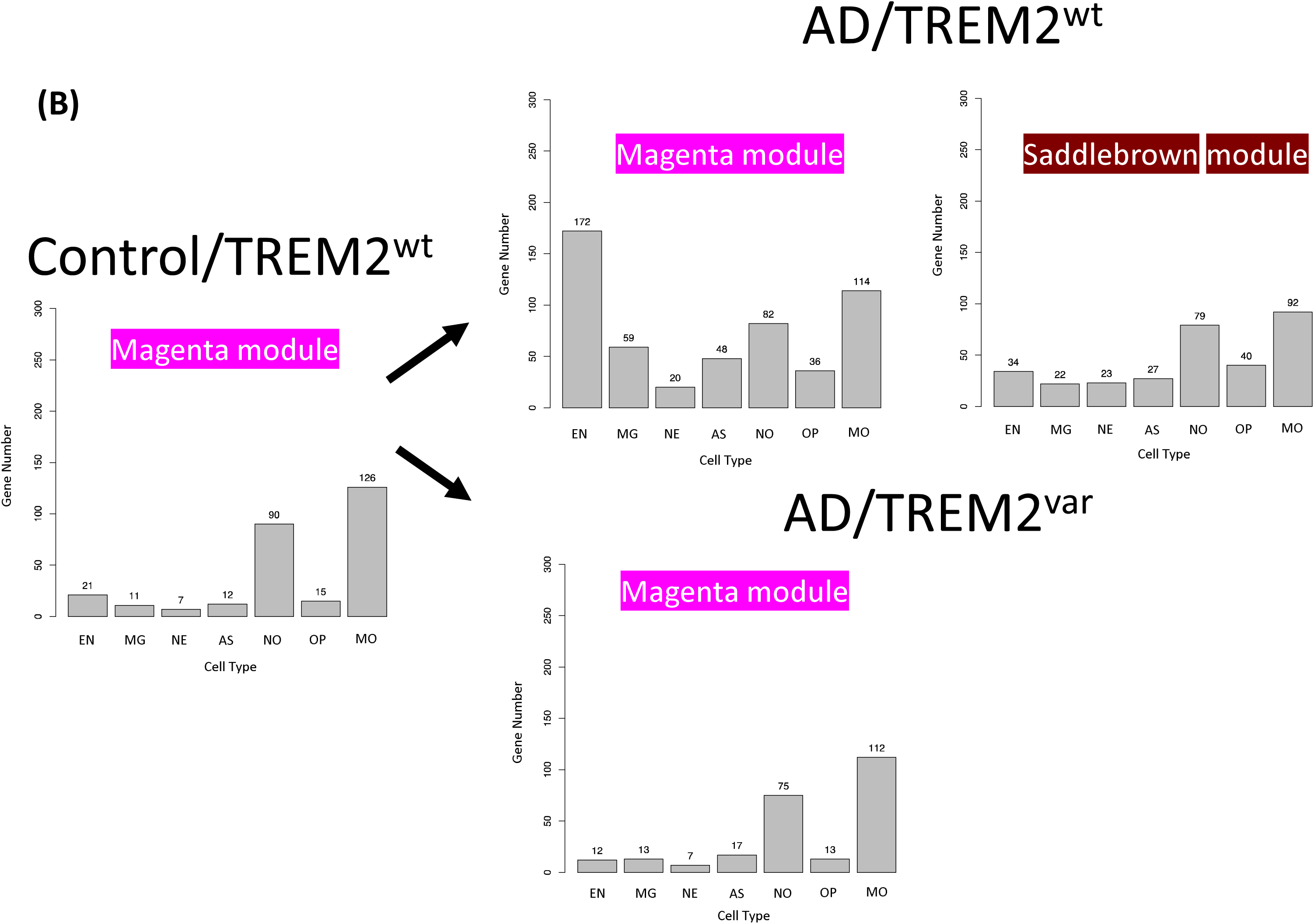
The cell type identity of the network modules was inferred using gene cell marker enrichment. Impact of AD and TREM2 variants in **Control/TREM2^wt^**, **AD/TREM2^wt^** and **AD/TREM2^var^** in key microglia and oligodendrocyte modules are presented: **(A)** Impact on modules with microglia identity. TREM2 and TYROBP were highly correlated with the purple module in **Control/TREM2^wt^** and **AD/TREM2^wt^**. TREM2 and TYROBP completely lost their central influence in **AD/TREM2^var^** and the module fragmented to become the blue and saddlebrown modules **(B)** Impact on modules with newly formed or myelinating oligodendrocyte identity. The magenta module with enrichment of genes from newly formed or myelinating oligodendrocytes split in to two modules in **AD/TREM2^var^** with the magenta module becoming highly correlated with a number of endothelial marker genes. This did not occur in **AD/TREM2^var.^** The magenta module instead remained similar to that observed for **Control/TREM2^wt^**. Each panel displays the proportion of markers for each cell type present in the module. The counts of the number of markers per cell type was divided by the number of markers present for that given cell type to take into account possible bias (see Methods for further details). The method allows us to approximate the proportion of each cell type represented in a given module. Key to cell types: endothelial cells (EN), microglia (MG), neurons (NE), astrocytes (AS), newly formed oligodendrocytes (NO), oligodendrocyte precursor cells (OP), myelinating oligodendrocytes (MO).

At the same time, a number of genes in the magenta module in **Control/TREM2^w^**^t^, which contained many genes enriched in newly formed and myelinating oligodendrocytes (Figure 5B), either remained in the magenta oligodendrocyte module which additionally took on genes highly enriched in endothelial cells, or, joined the saddlebrown module in **AD/TREM2^wt^** which retained a slightly reduced but still distinctive oligodendrocyte identity (Figure 5B). In contrast, the magenta module in **AD/TREM2^var^** remained largely intact and very similar to that found in people with no AD pathology (Figure 5B). Thus overall, the close relationship in expression between microglia, oligodendrocyte and endothelial genes observed in AD failed to fully express itself in those with TREM2 dysfunction where there was a much more limited correlation involving microglia and oligodendrocyte, but not endothelial modules.

The change in relationships between modules (and cell types) across the networks identified in **Control/TREM2^wt^**, **AD/TREM2^wt^** and **AD/TREM2^var^** was visualised through scaled 3D graphical representations using Cytoscape (Figure 6A). The microglia, endothelial and oligodendrocyte modules observed in **AD/TREM2^wt^** appeared to strongly rely on TREM2 and TYROBP which directly connected complement and ribosomal subunit genes within the purple microglia module (Figure 6B). Without their influence, the microglia-endothelial relationship was completely lost in **AD/TREM2^var^** and the connection between microglia and oligodendrocytes was weakened (Figure 6).

**Figure 6:**
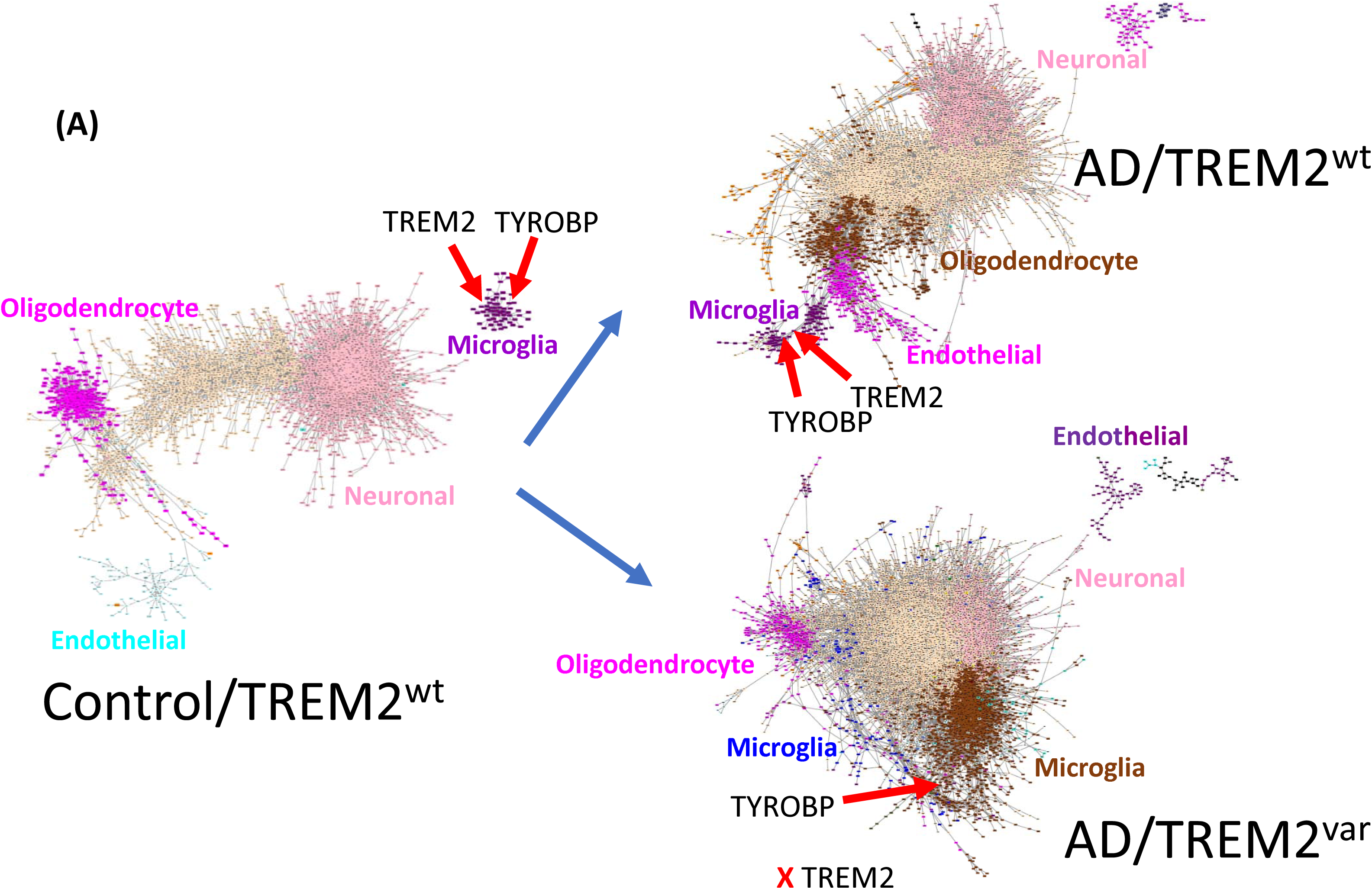
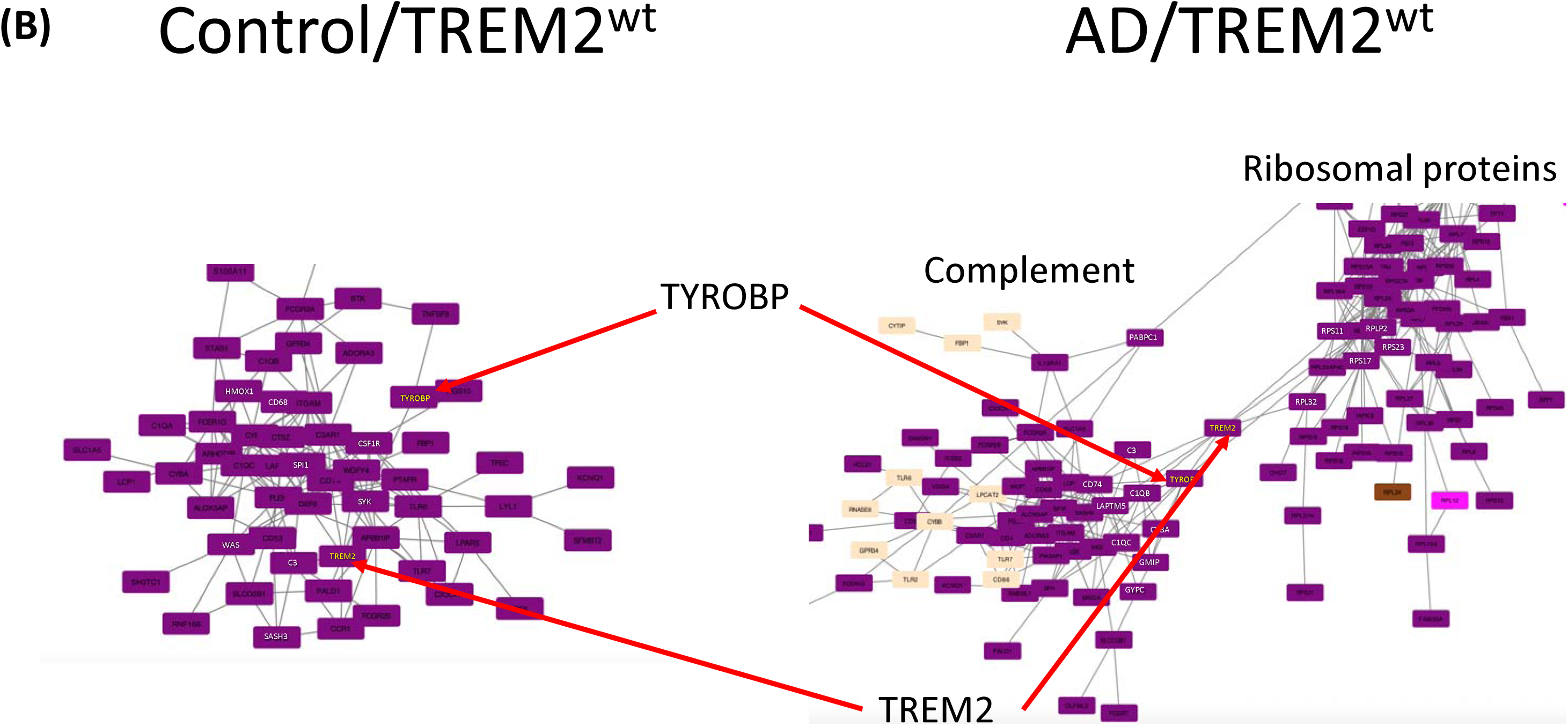
Graphical representation of the relationships between modules and how they change between **Control/TREM2^wt^**, **AD/TREM2^wt^** and **AD/TREM2^var^** in hippocampus. The network maps presented are the most highly correlated genes and modules for each group, retaining the same colours used within their respective group and matching the ones generated in WGCNA. We only display the genes with correlations stronger than 0.5 for clarity. **(A)** The whole network for each group, with individual modules labelled to indicate their cell type identity. **(B)** A closeup of the purple module with microglia identity in **Control/TREM2^wt^** and **AD/TREM2^wt^** highlighting the re-organisation of genes within this module in AD coinciding with a rise in prominence of TREM2 and TYROBP in **AD/TREM2^wt^**, which emerge to directly connect complement and ribosomal subunit genes within this module. TREM2 and TYROBP completely lose this influence in **AD/TREM2^var^** retaining only a weak correlation within the saddlebrown module which has a far weaker microglia identity.

### TREM2 and TYROBP impact ribosomal subunit genes in microglia and oligodendrocytes

Gene ontology and pathway analysis were used to reveal further biological insights in to the gene membership of each module and thereby further reveal the impact of AD and dependency of TREM2 on their behaviour. Significant gene enrichment for a number of biological pathways were found in modules particularly susceptible to AD and TREM2 dysfunction (Table S12-S21). Notably, the purple-microglia module in **AD/TREM2^wt^** (Table S7) was significantly enriched in “Ribosome, hsa03010, KEGG” genes (adj. p=5×10^-64^), largely consisting of genes coding for ribosome subunits (Table S12). These genes had previously resided in the non-microglial darkolivegreen module in **Control/TREM2^wt^** (Table S6) or became split across several modules in **AD/TREM2^var^** (Table S8), and were thus no longer collectively over-represented together as a group in any module.

As noted above, the magenta-endothelial module became much more closely aligned with oligodendrocyte genes and the purple-microglia module in **AD/TREM2^wt^** (Table S7) and in so doing, displayed moderate enrichment in “ECM-receptor interaction, hsa04512, KEGG” genes (adj. p=1×10^-7^) (Table S13). The equivalent endothelial module in **Control/TREM2^wt^** (Table S6, lightcyan) and **AD/TREM2^var^** (Table S8, purple and darkmagenta) displayed quite different biological profiles, namely, “Proteoglycans in cancer, hsa05205, KEGG” genes, adj. p=9×10^-5^, Table S14, “Proteoglycans in cancer, hsa05205, KEGG” genes, adj. p=1×10^-5^, Table S15 and “Hippo signaling pathway, hsa04390 KEGG” genes, adj. p=2×10^-3^, Table S16, respectively, and the ECM-receptor interaction genes were no longer enriched in these modules (Table S6 and S8).

The prominent saddlebrown-oligodendrocyte module in **AD/TREM2^wt^** (Table S7) had significant enrichment in “Proteasome, hsa03050, KEGG” genes (adj. p=2×10^-8^) (Table S17). In contrast, the equivalent oligodendrocyte modules in **Control/TREM2^wt^** (Table S6, magenta) and **AD/TREM2^var^** (Table S8, magenta) had no significant biological identity (Table S18 and S19). Interestingly, the same proteosome genes either resided together in the large pink-neuronal module in **Control/TREM2^wt^** or the saddlebrown-microglia module in **AD/TREM2^var^**, and for this reason were also significantly enriched as a group in these modules (“Proteasome, hsa03050, KEGG” genes, adj. p=1×10^-5^, Table S20, pink-neuronal module in **Control/TREM2^wt^** and “Proteasome, hsa03050, KEGG” genes, adj. p=1×10^-12^, Table S21, saddlebrown-microglia module in **AD/TREM2^var^**).

### Oligodendrocyte modules had the greatest proportion of TREM2-dependent DE genes compared DE genes commonly affected by AD

The modules to which the DE genes were assigned was re-examined to further explore the impacts of TREM2 on AD pathology. Importantly, we wanted to reveal genes, pathways and cell types in AD directly dependent on TREM2 (Figure 7). We used the used the top 1000 ranked genes in each DE comparison to classify them as either <u>AD changes independent of TREM2</u> (intersection of **Control/TREM2^wt^** and **AD/TREM2^wt^** DE genes *and* **Control/TREM2^wt^** and **AD/TREM2^var^** DE genes), <u>AD changes dependent on TREM2</u> (**Control/TREM2^wt^** and **AD/TREM2^wt^** DE genes *not in* **Control/TREM2^wt^** and **AD/TREM2^var^** DE genes) or <u>AD independent but TREM2 dependent changes</u> (**Control/TREM2^wt^** and **AD/TREM2^var^** DE genes *not in* **Control/TREM2^wt^** and **AD/TREM2^wt^** DE genes) (Figure 7A and Tables S2-5).

**Figure 7:**
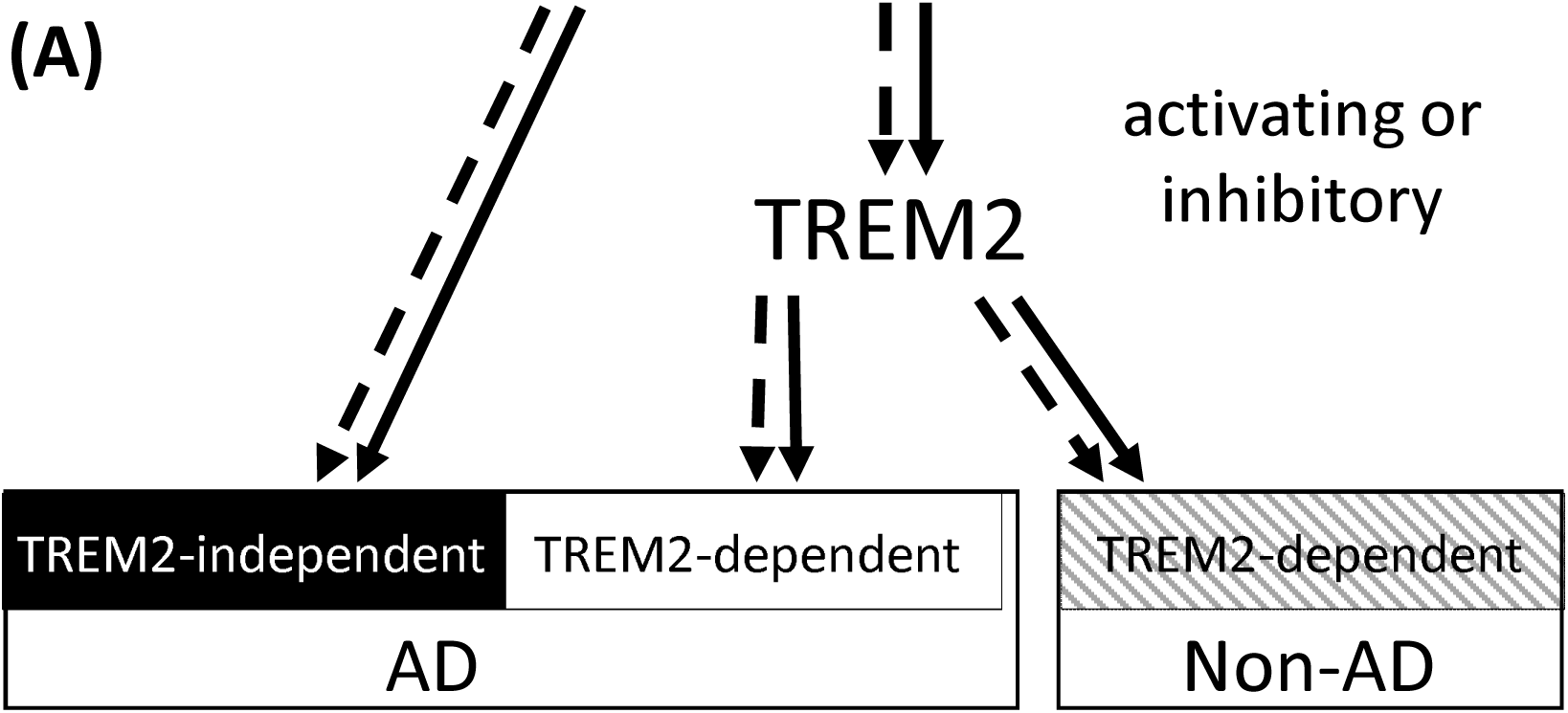
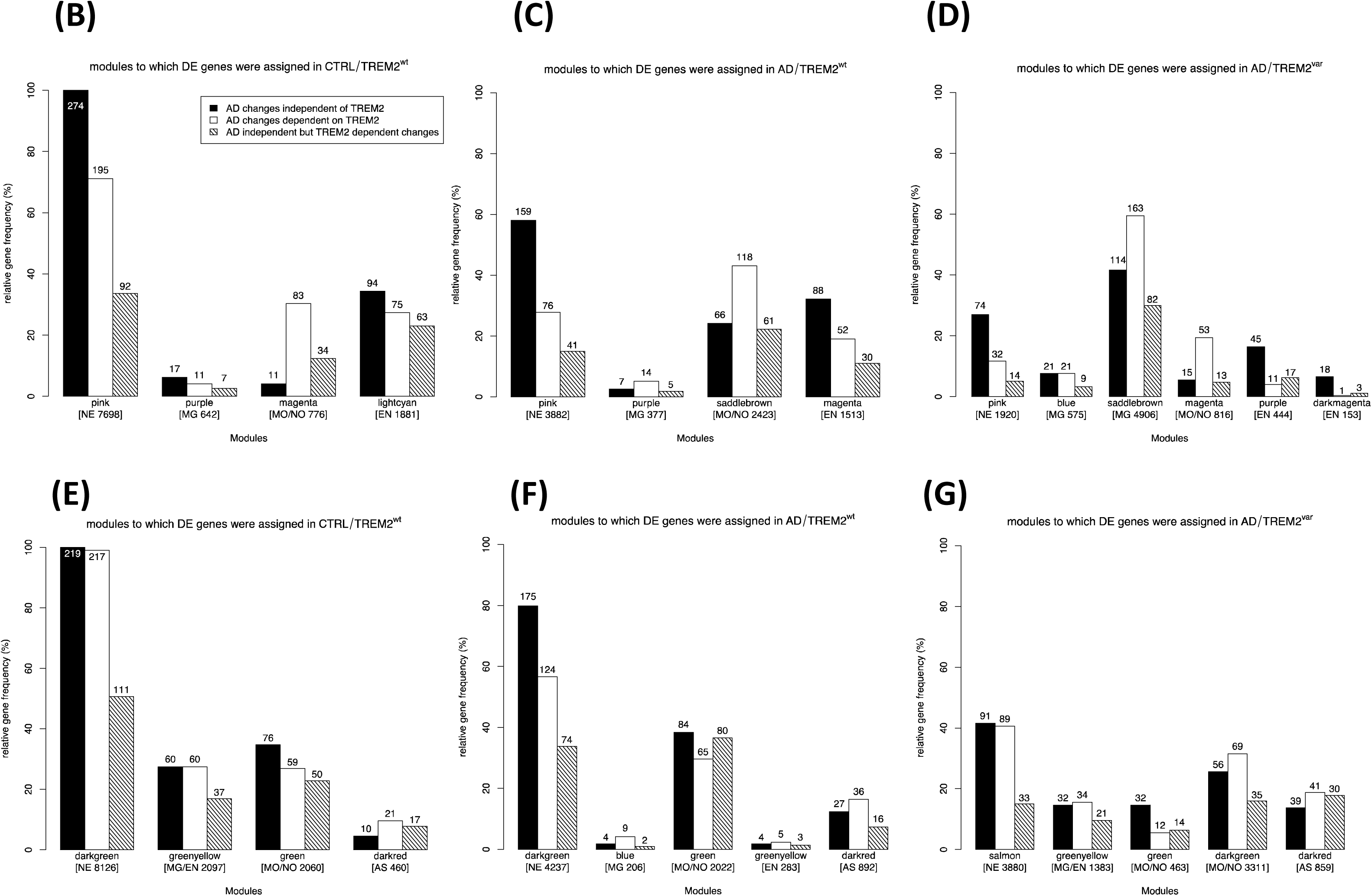
Evaluation of modules to which DE genes were assigned in hippocampus **(B-D)** and BA9 **(E-G)** and the impact of TREM2 and AD pathology **(A)** on these modules in **Control/TREM2^wt^ (B & E)**, **AD/TREM2^wt^ (C & F)** and **AD/TREM2^var^ (D & G)** brain samples. The 1000 top ranking by p-value DE genes were classified as either <u>AD changes independent of TREM2</u> (intersection of **Control/TREM2^wt^** and **AD/TREM2^wt^** DE genes and **Control/TREM2^wt^** and **AD/TREM2^var^** DE genes, black bars), <u>AD changes dependent on TREM2</u> (**Control/TREM2^wt^** and **AD/TREM2^wt^** DE genes *not in* **Control/TREM2^wt^** and **AD/TREM2^var^** DE genes, white bars) or <u>AD independent but TREM2 dependent changes</u> (**Control/TREM2^wt^** and **AD/TREM2^var^** DE genes *not in* **Control/TREM2^wt^** and **AD/TREM2^wt^** DE genes, hatched bars). Each bar represents the relative percentage (and number) of DE genes in each module and group **(B-G)**.

As expected, in hippocampus and BA9 the pink “neuronal” modules had the most DE genes across all groups of which DE genes common to AD dominated, rather than those dependent on TREM2. Although there was still an appreciable number of TREM2-dependent genes with altered expression residing in these modules, overall they were similarly represented across the groups and alterations in their expression didn’t appear to dramatically disrupt the patterning of neuronal modules (Figure 7B-D). Similarly AD dominated modules with endothelial identity rather than those dependent on TREM2. However unlike neuronal modules the proportion of TREM2- dependent genes residing in the magenta module in **AD/TREM2^wt^** hippocampus was markedly reduced compared to **Control/TREM2^wt^** and largely disappeared in the purple- and dark magenta-endothelial modules in **AD/TREM2^var^**, suggesting whilst TREM2 does’t affect the expression of all endothelial genes, it does appear important for maintaining some cohesiveness within the endothelial module in AD and thereby its relationship with other cell types in AD.

Few DE genes were assigned to microglia modules in hippocampus or BA9. This is not suprising as microglia form a relatively small fraction in whole brain samples so only the most highly expressed genes will be captured. Nevertheless, those that were differentially expressed in the purple-microglia module in the hippocampus of **AD/TREM2^wt^** had a clear TREM2 dependency. And in **AD/TREM2^var^** almost no DE genes were found in the blue- or saddlebrown-miroglia modules, further highlighting the central position of TREM2 in microglia in AD and the impact of losing TREM2 influence in microglia.

Modules with oligodendrocyte identity in hippocampus were also dominated by DE genes dependent on TREM2, suggesting, TREM2 (and microglia) have a central influence over oligodendrocyte function in AD. In BA9, this appeared to lead to the fragmentation of the oligodendrocyte module as many TREM2-dependent DE genes aligned with an additional new dark green-oligodendrocyte module in **AD/TREM2^var^**.

## Discussion

As we expected, overall the changes captured in the brain by whole genome gene expression analysis were overwhelmingly common to all AD samples, irrespective of TREM2 risk variant carrier status. However, we have also gained important insights in to additional TREM2- dependent changes in genes, pathways and cell types expected to significantly impact AD.

We find that microglia re-align their gene expression closely with that of oligodendrocytes and endothelial cells in the hippocampus of AD patients. This microglia-oligodendrocyte-endothelial expression re-alignment was dependent on TREM2-expressing microglia and AD pathology, as no such relationship was seen in control or AD cases with a TREM2 risk variant. TREM2-expressing microglia are therefore expected to play a prominent role in myelin and vascular integrity, particularly under disease conditions. Healthy myelination is crucial for the establishment and maintainenance of memories in the adult brain (Pan *et al*., 2020; Wang *et al*., 2020b). This capacity diminishes with age and disease. Significant myelin damage and oligodencrocyte dysfunction are a recognised feature of AD, even preceding clinical diagnosis (Gao *et al*., 2011; Benitez *et al*., 2014; Caso *et al*., 2015; Dean *et al*., 2017; Nasrabady *et al*., 2018; Mathys *et al*., 2019), although the underlying disease mechanisms in AD and its relationship with cells such as microglia have not been fully explored. A mechanistic link is more established for Multiple Sclerosis, where cycles of myelin damage and repair involving microglia are directly connected to neurological symptoms (Compston and Coles, 2008). When TREM2 signalling is blocked in an EAE mouse model of Multiple Sclerosis, there is greater demyelination and a worse clinical course (Piccio *et al*., 2007). It is also noteworthy that myelin pathology and hypomyelinosis attributable to oligodendrocyte dysfunction is a prominent feature in Nasu-Hakola disease caused by homozygous TREM2 or TYROBP loss-of-function mutations, a disease where axonal spheroids, neurodegeneration and cognitive decline occur at the latter stages of disease, but where characteristic amyloid and Tau AD pathology are absent (Yokoi *et al*., 1988; Tanaka, 2000; Paloneva *et al*., 2001; Kaifu *et al*., 2003; Klunemann *et al*., 2005; Turnbull and Colonna, 2007; Kaneko *et al*., 2010). Together these data suggest that normal TREM2 function and microglia are essential to counteract pathologies likely to involve compromises to myelin which also implicates oligodendrocytes.

Much of the work on TREM2 in AD has focused on it’s role in amyloid-β and Tau pathology. Overall, boosting or restoring TREM2 is considered beneficial for these pathologies and improves clinical outcomes in AD (Schlepckow *et al*., 2020). Loss of TREM2 on the other hand can lead to altered amyloid-β plaque morphology and/or increased susceptibility to tau seeding and spread in dystrophic neurons surrounding amyloid-β plaques in familial models of AD (Ulrich *et al*., 2014; Jay *et al*., 2015; Wang *et al*., 2015; Wang *et al*., 2016; Yuan *et al*., 2016; Bemiller *et al*., 2017; Jay *et al*., 2017; Cheng-Hathaway *et al*., 2018; Sayed *et al*., 2018; Song *et al*., 2018; Leyns *et al*., 2019; Parhizkar *et al*., 2019). While microglia and TREM2 can clearly impact these AD pathologies, our data does not capture any obvious TREM2-dependent expression profiles pointing to genes or pathways of influence in the AD hippocampal or cortical samples we examined that specifically impact these pathologies. This may be because the microglia changes we observe represent a common microglia response to a wide range of AD pathologies (amyloid, Tau, myelin and cell damage). Certainly, the expression profile of microglia surrounding amyloid (DAM-microglia) (Keren-Shaul *et al*., 2017) ovelaps with that found in white matter (PAM-microglia) associated with developmental remodelling (Li *et al*., 2019) despite each involving different cell-types and processes. Interestingly, PAM-microglia were found to phagocytose newly formed oligodendrocytes, alhough this was considered TREM2-independent (Li *et al*., 2019). In fact, around half of the genes we found in the purple-microglia module in AD are highly enriched in DAM microglia, notably TREM2, TYROBP, CD74, CD68 and both subunits of the complement receptor CR3 receptor (Mac-1; ITGAM/ITGB2; CD11b/CD18) and have previously been found to have correlated expression in AD (Zhang *et al*., 2013; Haure-Mirande *et al*., 2019), normal brain (Hawrylycz *et al*., 2012; Forabosco *et al*., 2013; McKenzie *et al*., 2018; Olah *et al*., 2018) and a TREM2 KO mouse model (Carbajosa *et al*., 2018). However, we also found additional AD-associated microglia genes in our study which were not prominently enriched in DAM, namely impacts on ribosomal subunit genes in microglia and proteosomal subunit genes in oligodendrocytes (discussed below). As DAM cells are a relatively small proportion of the total microglia population (Keren-Shaul *et al*., 2017), it is possible that the amyloid/Tau responding microglia are harder to detect in our whole brain samples, although we could readily detect the majority of microglia-specific genes previously reported (Zhang *et al*., 2014). We could also readily detect genes highly enriched or unique to astrocytes, but unlike the other major cell types present in brain, somewhat surprisingly their expression was not sufficiently co-ordinated to form a unique module of their own in our data which would allow us to explore their behaviour towards other modules or cell types in AD brain and in relation to TREM2.

Microglia are capable of phagocytosing a range of endogenous and pathogenic cargoes in response to different DAMPs and PAMPs from whole cells to ‘nibbling’ cell membranes such as occurs in synapse remodelling (Schafer *et al*., 2012; Hong *et al*., 2016; Vasek *et al*., 2016; Weinhard *et al*., 2018; Wang *et al*., 2020a). Loss of TREM2 signalling adversely impacts phagocytosis (Takahashi *et al*., 2005; Hsieh *et al*., 2009; N’Diaye *et al*., 2009; Stefano *et al*., 2009; Kleinberger *et al*., 2014; Atagi *et al*., 2015; Gawish *et al*., 2015; Kawabori *et al*., 2015) while boosting TREM2 signalling increases phagocytosis (Takahashi *et al*., 2005; Takahashi *et al*., 2007; Jiang *et al*., 2014; Kleinberger *et al*., 2014). TREM2 appears to be a DAMP receptor exclusively expressed by microglia in the brain able to respond to anionic and zwitterionic lipids, including sphingomyelin, phosphatidylserine (PS) and phosphatidylethanolamine (PE) (Daws *et al*., 2003; Cannon *et al*., 2012; Wang *et al*., 2015; Shirotani *et al*., 2019). These are major components of both myelin and cell membranes (O’Brien and Sampson, 1965; Aureli *et al*., 2015), with PS and PE exposure on degenerating cells marking them out for microglia clearance (Nagata *et al*., 2016; Shirotani *et al*., 2019). TREM2 has previously been shown to have a role in clearance of myelin debris (Poliani *et al*., 2015) engulfment of apoptotic neurons (Takahashi *et al*., 2005; Hsieh *et al*., 2009; Garcia-Reitboeck *et al*., 2018; Shirotani *et al*., 2019) and lysosomal degradation and egress of cholesterol and lipid products (Nugent *et al*., 2020). Microglia can also clear oligodendrocytes (Li *et al*., 2019), although it is unclear whether this is a TREM2-dependent process in the context of AD.

One of the most striking findings in our data is the co-ordinated expression of an FcγR- complement mediated phagocytic microglia response in AD which relies on TREM2 function. It was highly correlated and reliant on TREM2-DAP12 for its correlation and connection with ribosomal genes in the microglia module and proteosomal genes in the oligodendrocyte module. In addition to TREM2, TYRPOBP, CSFR1 and SPI1, amongst the most significantly correlated 109 microglia-purple module genes in AD, there were Fc receptor genes (FCGR1A/FCGR1B [FcγI; CD64], FCGR2A/FCGR2B [FcγII; CD32], FCGRT, FCER1G), classical complement pathway activation genes (C1QA, B & C, C3, C3AR1, VSIG4 and ITGB2/ITGAM [CR3; Mac-1; CD11b/CD18; αMβ2]), phagocytic/lysosomal markers (CD68, CYBA, LAPTM5, RAB3IL1), MHC Class II-antigen presentation genes (CD74, CD4, WDFY4, HCLS1) and 48 large and small ribosomal subunit genes. There were also many genes linked to the cytoskeletal system (ARHGDIB, APBB1IP, FGD2, GMIP, LCP1, SAMSN1) which presumably coordinates what appears to represent an active response to AD pathology. The coordinated expression of many of these genes in AD has been noted previously (Zhang *et al*., 2013; Matarin *et al*., 2015) and FCER1G, LAPTM5 and HCK, which are highly correlated and central to the AD purple-microglia module in our data have recently been identified as key drivers of an AD expression module role with effects on phagocytosis and lysosomal function in microglial cells (Patel *et al*., 2020). Together, these data suggest the TREM2 pathway is controlling an antibody-dependent cell mediated phagocytosis/cytotoxicity (ADCP/ADCC) and/or complement-dependent cell-mediated phagocytosis/cytotoxicity (CDCP/CDCC) in AD. We are unable to distinguish from our results whether ADCP/ADCC and CDCP/CDCC are both active in AD, are acting independently, synergistically or are protective or detrimental in AD, but oligodendrocytes and myelin appear to be the main target.

The coordinated expression of this module, particularly the genes detailed above, was disrupted in AD with a TREM2 risk variant. The expression of the Fc receptor genes, notably FCGR1B and FCGR2B, and the complement gene C1QA particularly had expression profiles which deviated from the other genes in the module suggesting TREM2 specifically impacts ADCP/ADCC and CDCP/CDCC and the humoral immune system. Evidence confirms many of these genes are uniquely expressed by microglia in the brain (McKenzie *et al*., 2018) and there are known functional links between them and TREM2. Fcγ receptors and TREM2 (via adapter DAP12; TYROBP) separately or cooperatively utilise ITAM and downstream signalling through Syk (Zou *et al*., 2008; Mócsai *et al*., 2010; Satoh *et al*., 2012b; Yao *et al*., 2019). DAP12 is also an adaptor for CR3 in microglia-mediated apoptic neuron clearance (Wakselman *et al*., 2008), highlighting the close functional relationship between Fc and complement. Knocking out DAP12 in a Tau (Audrain *et al*., 2019) or amyloid (Haure-Mirande *et al*., 2019) mouse model lowers DAM-associated genes including C1Q, and at least in the Tau model, led to an acceleration of Tau pathology. Somewhat surprisingly this was associated with clinical improvement in both mouse models (Audrain *et al*., 2019; Haure-Mirande *et al*., 2019) which is at odds with the clinical course in Nasu-Hakola disease. Others show loss of complement function leads to poorer outcomes in AD (Wyss-Coray *et al*., 2002; Maier *et al*., 2008) and mutations (particularly in C1QA) linked to SLE result in greater autoantibody production and loss of immunotolerance (Sontheimer *et al*., 2005). Syk phosphorylation is increased in Nasu-Hakola disease when TREM2 is absent (Satoh *et al*., 2012a) and heightened phopho Syk is also associated with autoimmune diseases such as Rheumatoid Arthirits (Mócsai *et al*., 2010). We speculate that compromised TREM2 in the presence of AD pathologies in microglia alterns the balance of Fcγ receptor activity and/or the generation of autoantibodies and/or the threshold required to respond to them, with impacts on the balance of ‘silent’ immune complement mediated phagocytosis and a pro-inflammatory response involving Fcγ receptors. A lack of timely clearance of exposed damage signals prevalent in AD, can lead to destructive self-antigens being generated as the immune pivots its activities against healthy cell or tissue targets (Park and Kim, 2017). Future work needs to explore these mechanisms further, particularly TREM2 dependency for Fcγ receptor mediated activity in AD, and its synergy with complement activation.

Our AD findings are consistent with a model of heightened pro-inflammatory response perpetuated through a chronic cycle inducing bystander damage, cell death and a shift away from immunotolerance previously described for AD. There is numerous evidence implicating immune activation in AD including inflammasome activation, pro-inflammatory cytokine release (Heneka *et al*., 2015), activation of complement intermediates such as the anaphylatoxins C3a and C5a (Eikelenboom and Veerhuis, 1996; Veerhuis, 2011; Morgan, 2018) and greater antigen presentation by MHC Class II (McGeer *et al*., 1987; Styren *et al*., 1990; Perlmutter *et al*., 1992). Complement activation is a well recognised response in AD brain (Hong *et al*., 2016; Morgan, 2018) and is a normal developmental mechanism for sculpting neural circuits (Schafer *et al*., 2012), but what has not previously been recognised is the central position the TREM2 pathway may play in this response and the close connection to oligodendrocytes. and by extension, an impact on myelin integrity.

Ribosomes consist of a large 60S subunit containing ∼30 “RPL” proteins and a small 40S subunit consisting of ∼20 “RPS” proteins (Schmeing and Ramakrishnan, 2009). It is thereore no surprise that they need to be coordinately expressed to form equimolar amounts for a functioning ribosome (Mayer and Grummt, 2006). The strong correlation with each other in our data exemplifies this. In our brain samples we detected expression of 251 RPL and 151 RPS different genes. Ribosome composition differs between cell types. We appear to have identified the microglia-specific ribosomal genes since the same genes were consistently found together in the same module across samples, they were strongly associated with other microglia genes in AD and the number we detect in this module is consistent with the number of subunits required to form a functioning ribosome. Similarly, a large subset of proteosomal subunit genes with co-ordinated expression in an oligodendrocyte module were positively aligned with the ribosomal and other microglia genes detailed above. Interestingly, this included PSME1 and PSME4, two of the four components of the 11S activator proteosomal particle, which has been shown to specialise in processing antigens in response to INFγ (Wang and Maldonado, 2006). The coordinated expression of the ribosomal genes in microglia and the proteosomal genes linked to an oligodendcrotye module suggest a ramping up of protein expression in AD alongside the immune/phagocyic activation in microglia and appears to be associated with apoptosis and/or cell stress in oligodendrocytes, pehaps linked to myelin damage and a damage and/or antigen-mediated immune response by microglia.

Overall, our data supports a role for TREM2 in mediating oligodendrocyte and/or myelin clearance in AD which may be essential not only for preserving healthy tissue homeostasis but may also serve to minimise the persistence of antigenic peptides and lipids which may lead to detrimental pro-inflammatory sequelae. We hypothesise that not enough TREM2 activity either as a result of its dysfunction as pathology emerges or when it becomes overwhelmed by pathologies which interfere with its ability to adequately clear damaged cells or tissue creates an imbalance in favour of a more damaging pro-inflammatory response leading to aggressive damage to cells and tissue, with myelin being particularly vulnerable.

## Acknowledgements

We would like to thank the donors and their families for making tissue available for this study. This project is co-funded by the Eli Lilly and Company Lilly Research Award Program (LRAP) which also supports GC, KM, NL, DAC and MJON who are employees of Eli Lilly and Company Ltd, UK. JR and HW are employees of Eli Lilly and Company (Indianapolis, USA). RJBD and SJN are also part of NIHR Biomedical Research Centre at South London and Maudsley NHS Foundation Trust and King’s College London, UK and Farr Institute of Health Informatics Research, UCL Institute of Health Informatics, University College London, London, United Kingdom. This project has also received support from the Innovative Medicines Initiative 2 Joint Undertaking under grant agreement No 115976. This Joint Undertaking receives support from the European Union’s Horizon 2020 research and innovation programme and EFPIA. RJBD, SJN are also supported by the National Institute for Health Research (NIHR) University College London Hospitals Biomedical Research Centre, and by awards establishing the Farr Institute of Health Informatics Research at UCL Partners, from the Medical Research Council, Arthritis Research UK, British Heart Foundation, Cancer Research UK, Chief Scientist Office, Economic and Social Research Council, Engineering and Physical Sciences Research Council, National Institute for Health Research, National Institute for Social Care and Health Research, and Wellcome Trust (grant MR/K006584/1). AH holds a Medical Research Council (MRC) eMedLab Medical Bioinformatics Career Development Fellowship, funded from award MR/L016311/1.

## Supplementary Figure and Table legends

<u>Supplementary File 1:</u> Quality control metrics including sample RNA integrity number PCA plots; sex concordance between recorded sex versus Y chromosome expression; concordancy of direction of effect of the differential expressed genes in **AD/TREM2^var^** network compared to the **AD/TREM2^wt^** network; Scale free topology Network Assesement

<u>Table S1:</u> Characteristics of cases **(A)** and samples (**B)** used in the study, including outcomes of data and sample quality control checks and justification for sample exclusion from differential expression or WGCNA analyses.

<u>Table S2:</u> Differentially expressed transcripts in **Control/TREM2^wt^ vs AD/TREM2^wt^** hippocampus

<u>Table S3:</u> Differentially expressed transcripts in **Control/TREM2^wt^ vs AD/TREM2^var^** hippocampus

<u>Table S4:</u> Differentially expressed transcripts in **Control/TREM2^wt^ vs AD/TREM2^wt^** BA9

<u>Table S5:</u> Differentially expressed transcripts in **Control/TREM2^wt^ vs AD/TREM2^var^** BA9

<u>Table S6:</u> Module membership of transcripts in **Control/TREM2^wt^** hippocampus

<u>Table S7:</u> Module membership of transcripts in **AD/TREM2^wt^** hippocampus

<u>Table S8:</u> Module membership of transcripts in **AD/TREM2^var^** hippocampus

<u>Table S9:</u> Module membership of transcripts in **Control/TREM2^wt^** BA9

<u>Table S10:</u> Module membership of transcripts in **AD/TREM2^wt^** BA9

<u>Table S11:</u> Module membership of transcripts in **AD/TREM2^var^** BA9

<u>Table S12:</u> KEGG pathway enrichment of genes in purple-microglia module in **AD/TREM2^wt^** hippocampus

<u>Table S13:</u> KEGG pathway enrichment of genes in magenta-endothelial module in **AD/TREM2^wt^** hippocampus

<u>Table S14:</u> KEGG pathway enrichment of genes in lightcyan-endothelial module in **Control/TREM2^wt^** hippocampus

<u>Table S15:</u> KEGG pathway enrichment of genes in purple-endothelial module in **AD/TREM2^var^** hippocampus

<u>Table S16:</u> KEGG pathway enrichment of genes in darkmagenta-endothelial module in **AD/TREM2^var^** hippocampus

<u>Table S17:</u> KEGG pathway enrichment of genes in saddlebrown-oligodendrocyte module in **AD/TREM2^wt^** hippocampus

<u>Table S18:</u> KEGG pathway enrichment of genes in magenta-oligodendrocyte module in **Control/TREM2^wt^** hippocampus

<u>Table S19:</u> KEGG pathway enrichment of genes in magenta-oligodendrocyte module in **AD/TREM2^var^** hippocampus

<u>Table S20:</u> KEGG pathway enrichment of genes in pink-neuronal module in **Control/TREM2^wt^** hippocampus

<u>Table S21:</u> KEGG pathway enrichment of genes in saddlebrown-microglia module in **AD/TREM2^var^** hippocampus

<u>Figure S1:</u> WGCNA modules and their preservation in BA9 between **Control/TREM2^wt^** and **AD/TREM2^wt^** networks **(A)**, and **Control/TREM2^wt^** and **AD/TREM2^var^** networks **(B)**. Each row of the table corresponds to one **Control/TREM2^wt^** set-specific module (assigned a colour as indicated by the text label). Each column corresponds to one **AD/TREM2^wt^** or **AD/TREM2^var^** set-specific module, respectively. Numbers indicate the number of genes in each module which are also divided across the table to indicate the number of genes which intersect each pair of modules between the two groups. Red shading within the table is generated by calculating the – log(p) with p being the hypergeometric test p-value for the degree of overlap of each module pair. The stronger the red color, the more significant the overlap is. Most **Control/TREM2^wt^** set-specific modules have a corresponding **AD/TREM2^wt^** counterpart or involve a split or merger involving only two modules **(A)**. This is also largely true between **Control/TREM2^wt^** and **AD/TREM2^var^ (B)**. Notably, the TREM2 and TYROBP containing greenyellow module in **Control/TREM2^wt^** is also preserved in **AD/TREM2^var^** but fragments in to the TREM2 and TYROBP containing blue module in **AD/TREM2^wt^**. Other modules are similarly affected suggesting TREM2 and TYROBP have limited influence in **Control/TREM2^wt^** and **AD/TREM2^var^** where its activity is either not needed or absent due to dysfunction. In contrast TREM2 and TYROBP have significant influence in **AD/TREM2^wt^**. The grey module which contains unassigned genes is excluded from consideration.

<u>Figure S2:</u> Preservation of **Control/TREM2^w^** modules in **AD/TREM2^wt^** and **AD/TREM2^var^** hippocampus **(A-D)** and BA9 **(E-H)** networks. Images in **(A, C, E, G)** show the composite statistic median rank versus module size. The higher the median rank the less preserved the module is relative to other modules. Since median rank is based on the observed preservation statistics (as opposed to Z statistics or p-values), it is independent of module size. Images in **(B, D, F, H)** show the composite statistic Z summary. If Z summary >10 there is strong evidence that the module is preserved (Langfelder *et al*., 2011). If Z summary *∖*textless 2, there is no evidence that the module is preserved. Note that Z summary shows a strong dependence on module size. Colours match those assigned in WGCNA analysis (Figure 4).

