## Supplemental Figures S1 and S2 for "TREM2 impacts brain microglia, oligodendrocytes and endothelial co-expression modules revealing genes and pathways important in Alzheimer’s disease"

### Slide 1
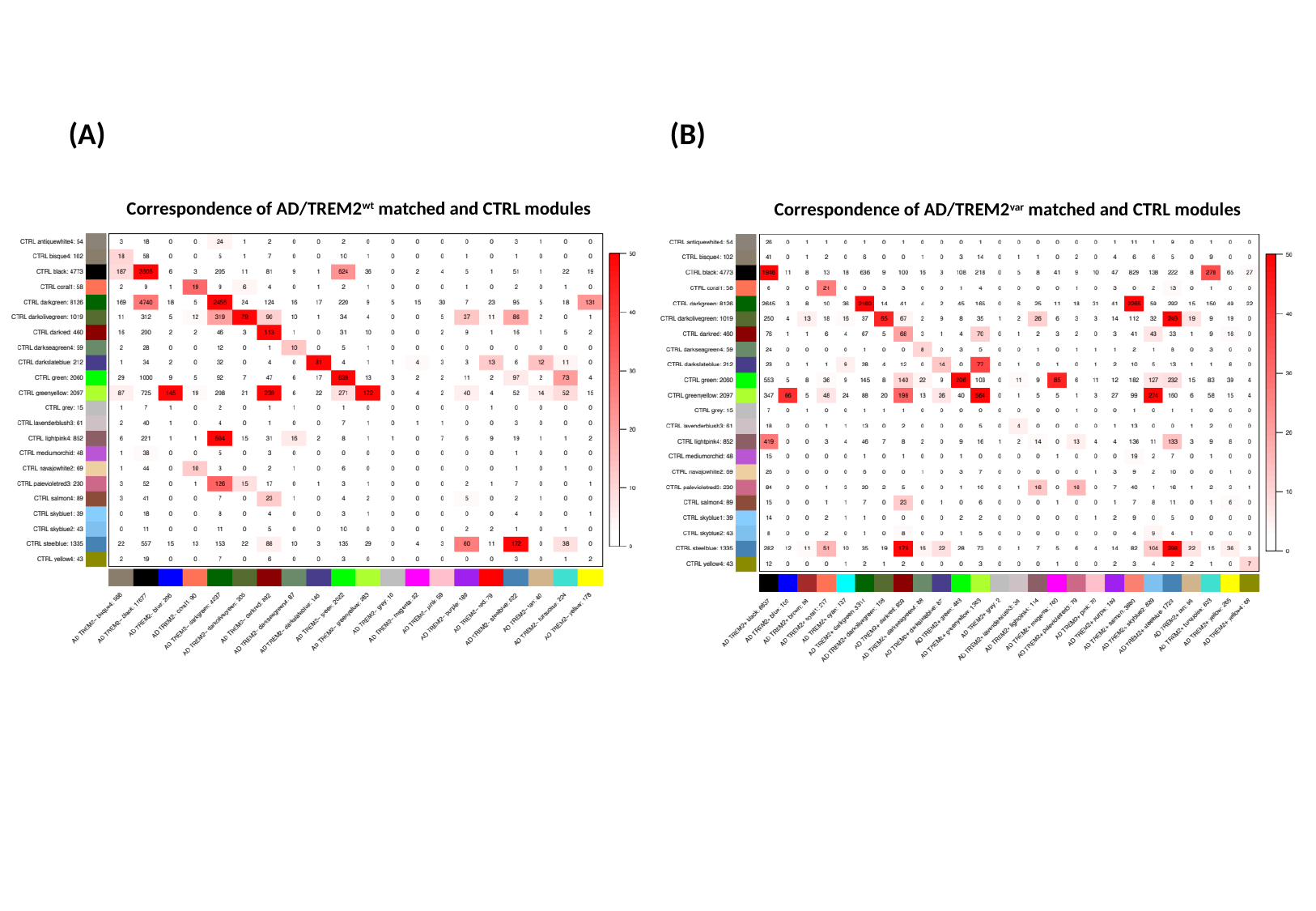

(A)
(B)
Correspondence of AD/TREM2^wt^ matched and CTRL modules
Correspondence of AD/TREM2^var^ matched and CTRL modules

### Slide 2
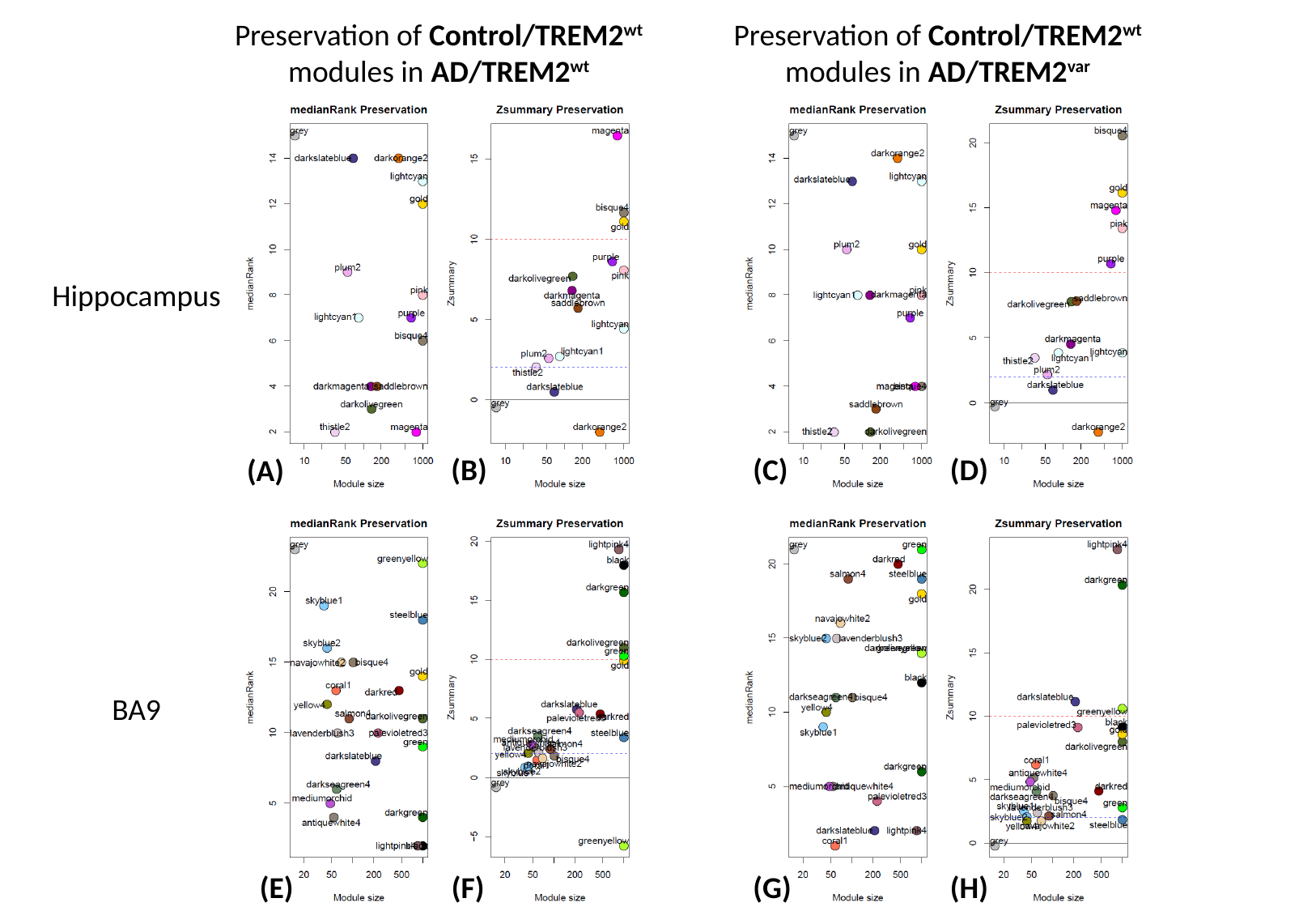

Preservation of Control/TREM2^wt^ modules in AD/TREM2^wt^
Preservation of Control/TREM2^wt^ modules in AD/TREM2^var^
Hippocampus
(B)
(C)
(D)
(A)
BA9
(E)
(F)
(G)
(H)
