## Supplemental File 1 for "TREM2 impacts brain microglia, oligodendrocytes and endothelial co-expression modules revealing genes and pathways important in Alzheimer’s disease"

PCA plot showing the samples clustering by RIN score.

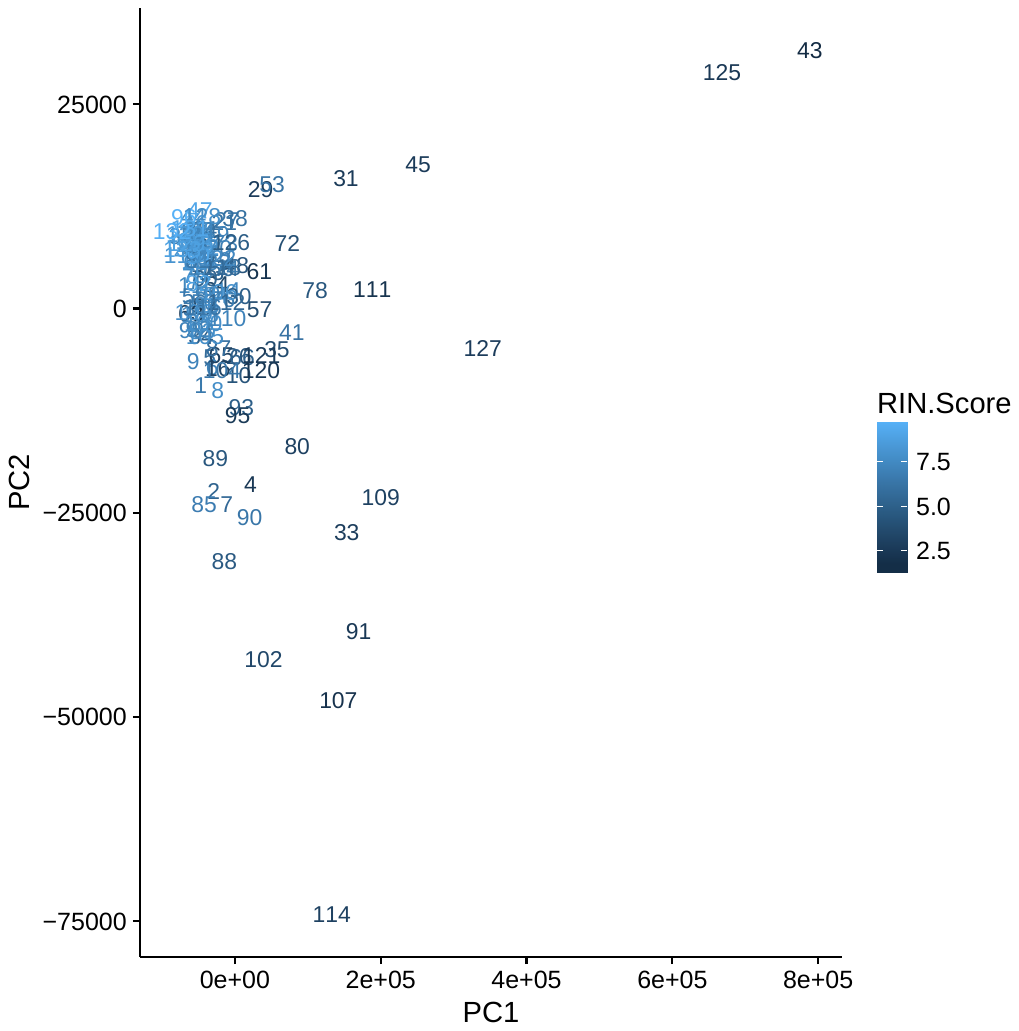

Number of variants Y variants called for each of the samples to confirm the validity of the sex annotations.
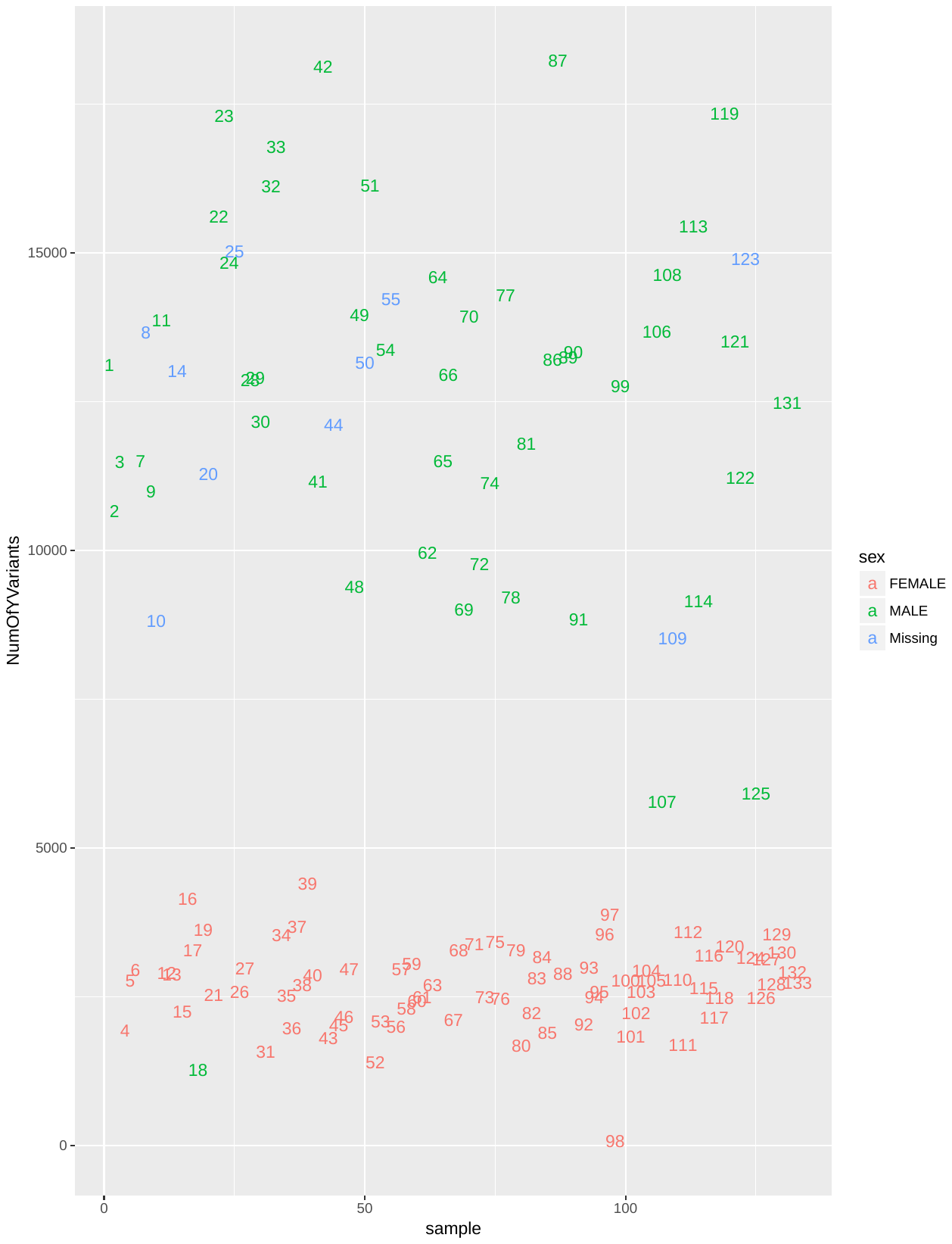

Concordancy of direction of effect of the differential expressed genes in **AD/TREM2^var^** network compared to the **AD/TREM2^wt^** network, showing the broad concordance, particularly for the top ranking genes.
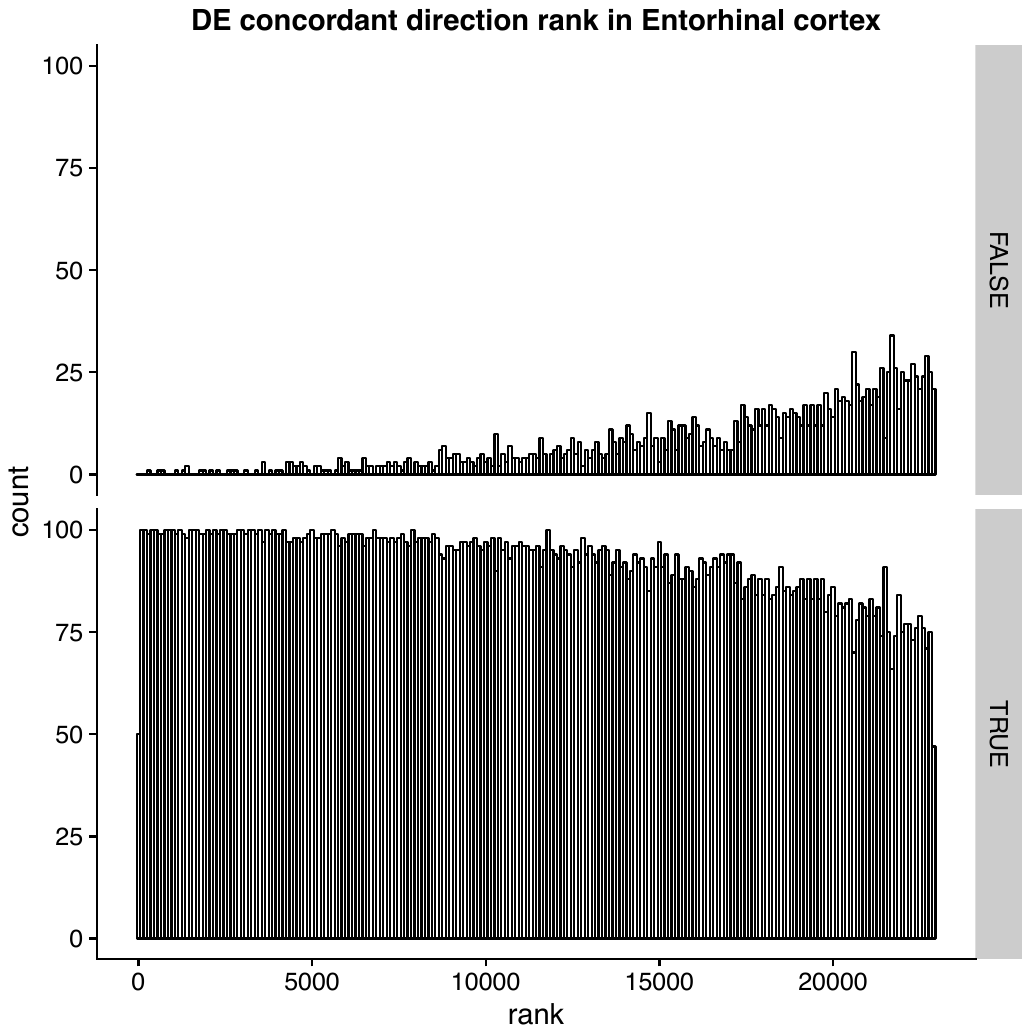

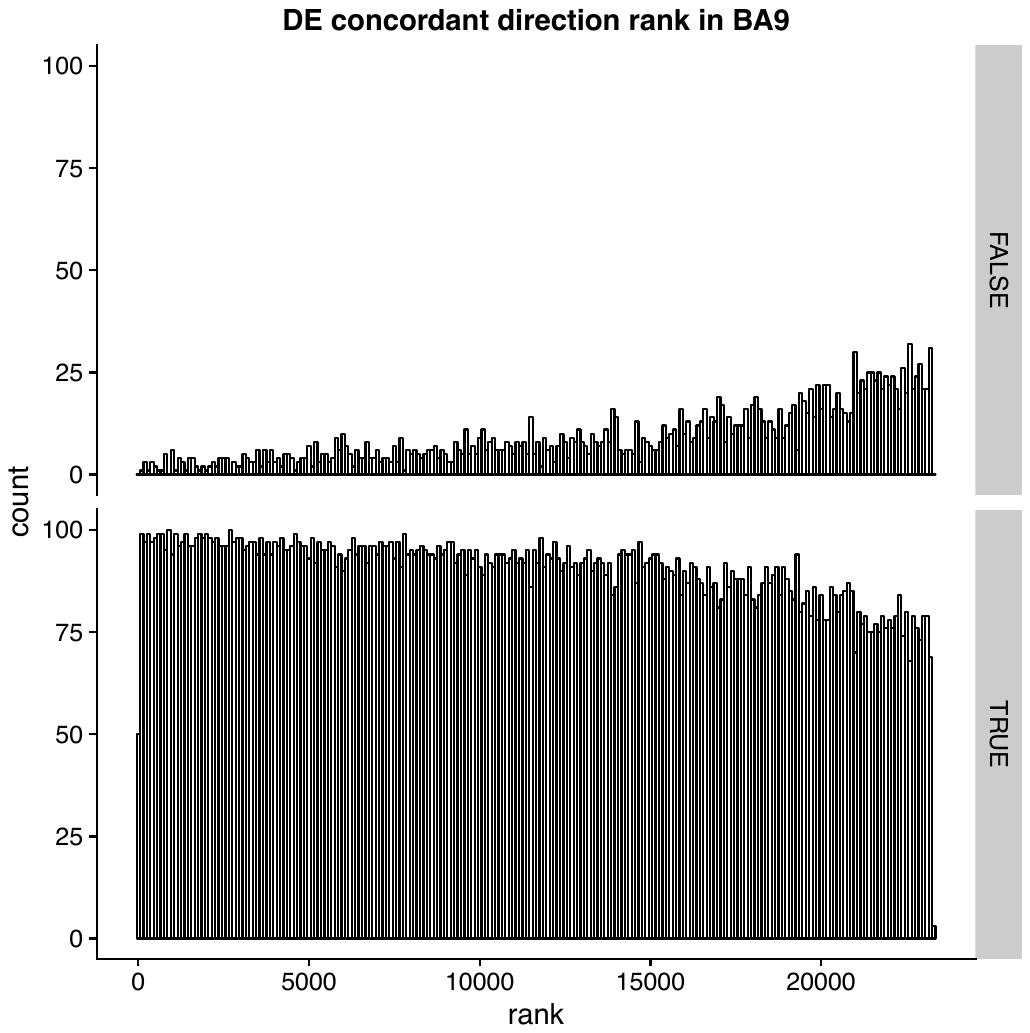
 Scale free topology Network Assesement in Entorhinal cortex: CTRL, AD and AD-TREM2 networks:
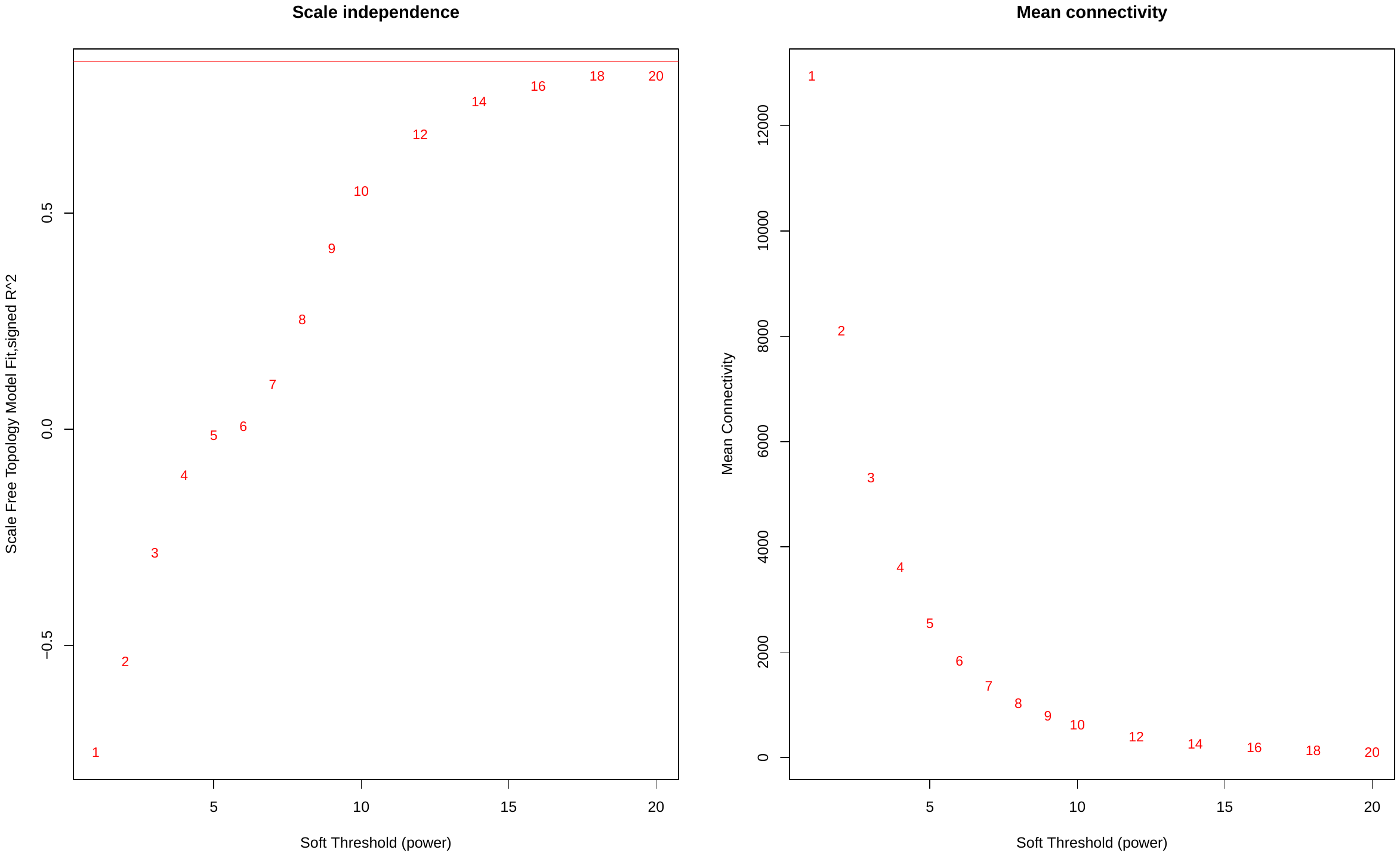

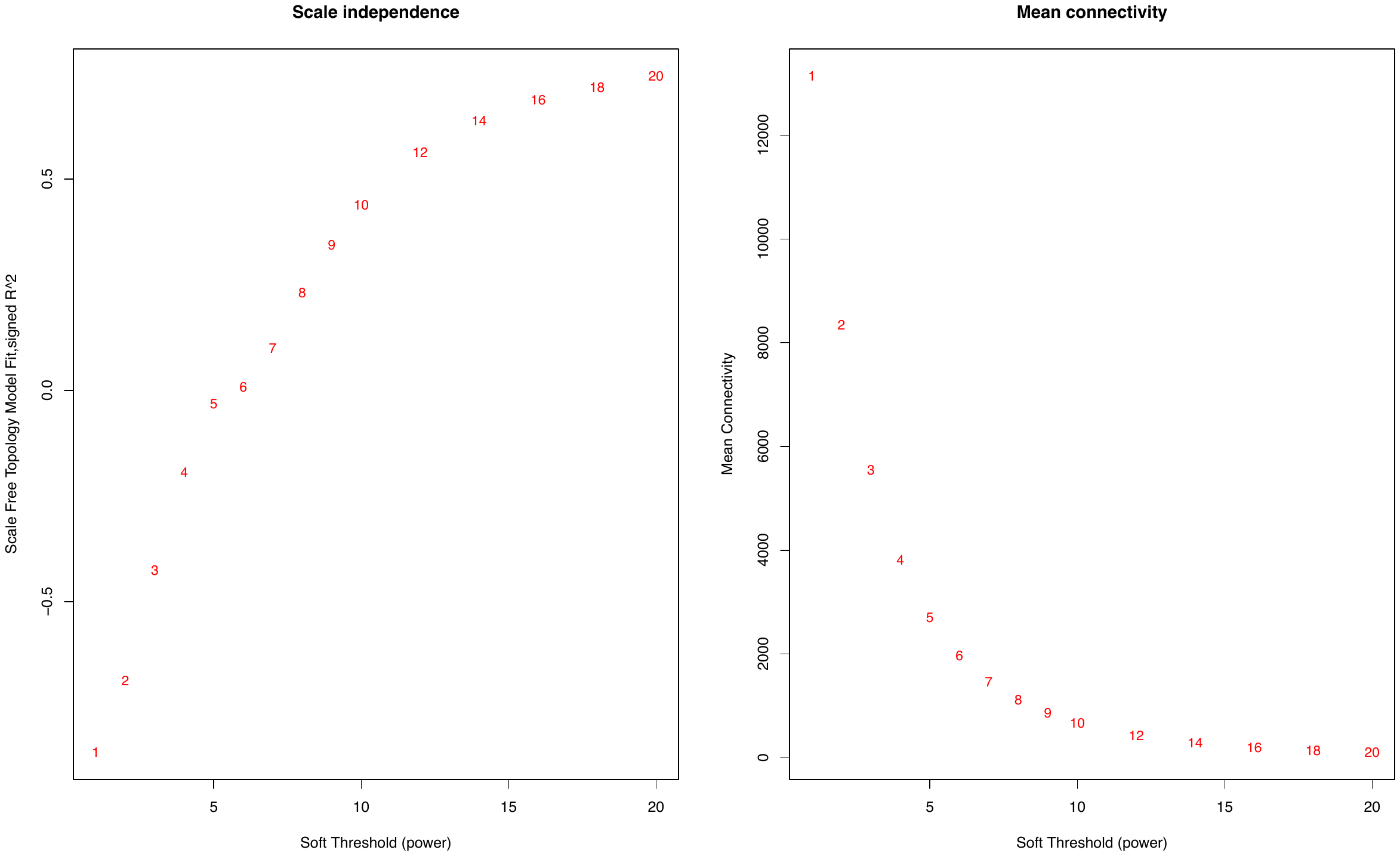

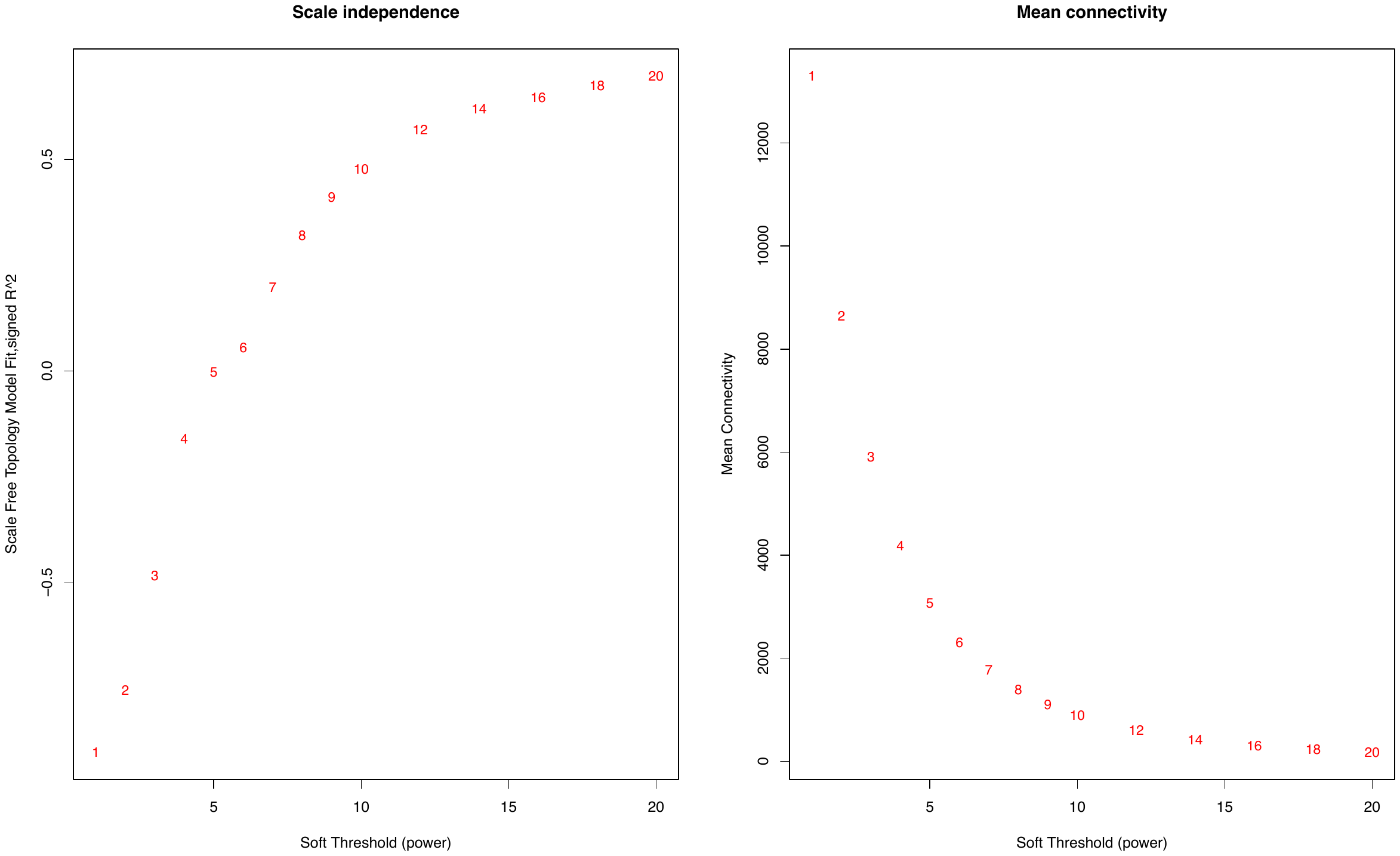
 Scale free topology Network Assesement in BA9: CTRL, AD and AD-TREM2 networks:

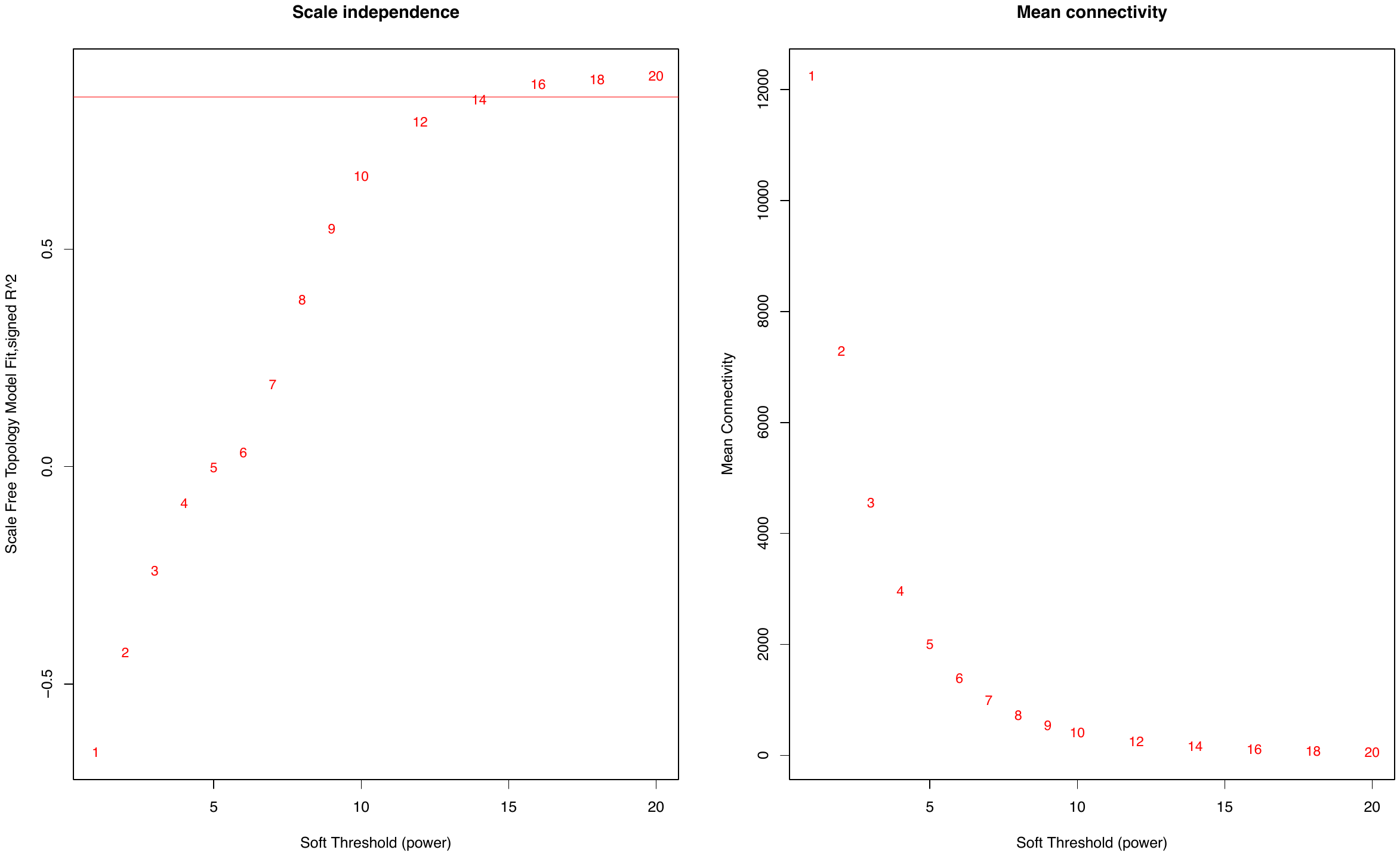

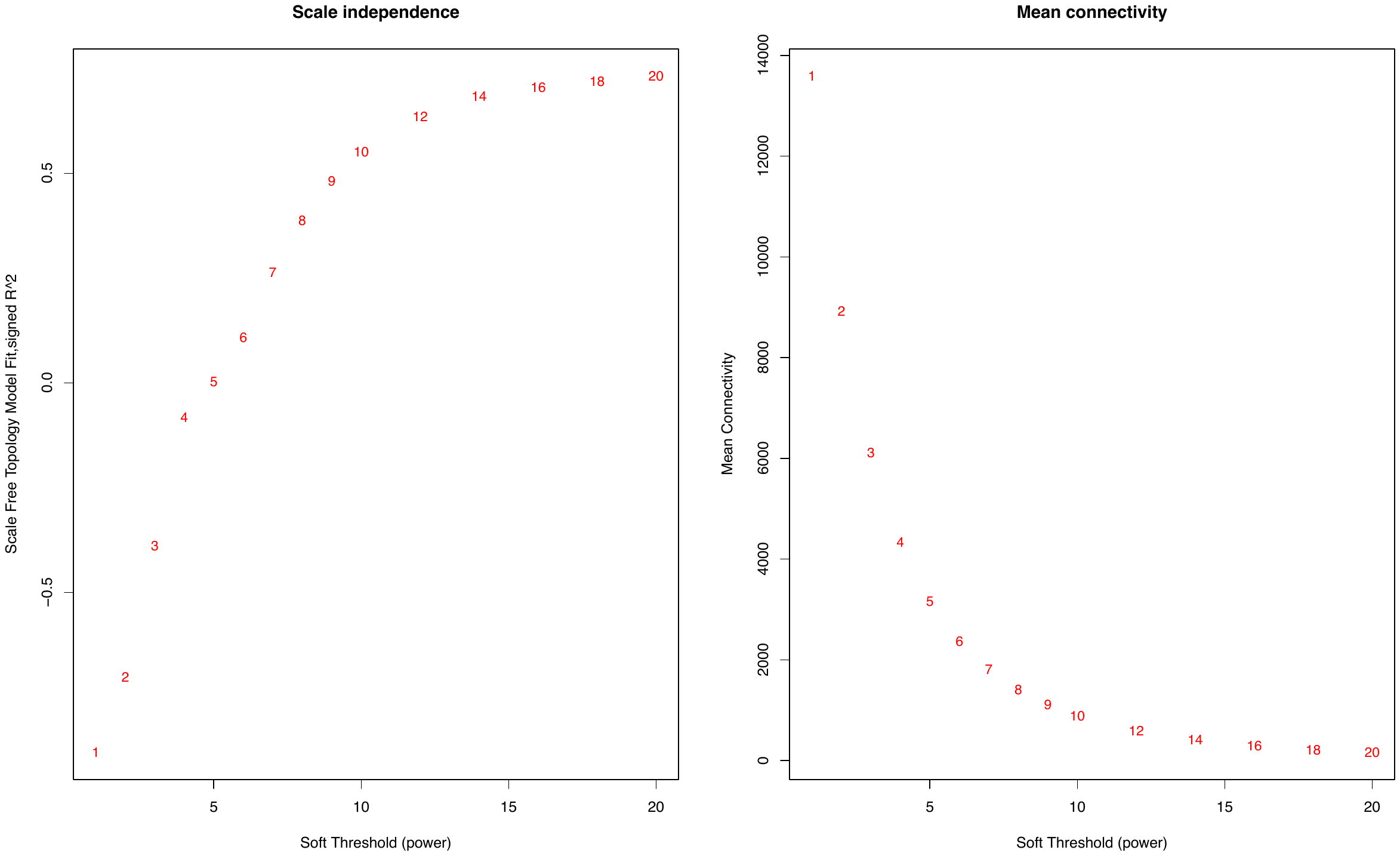

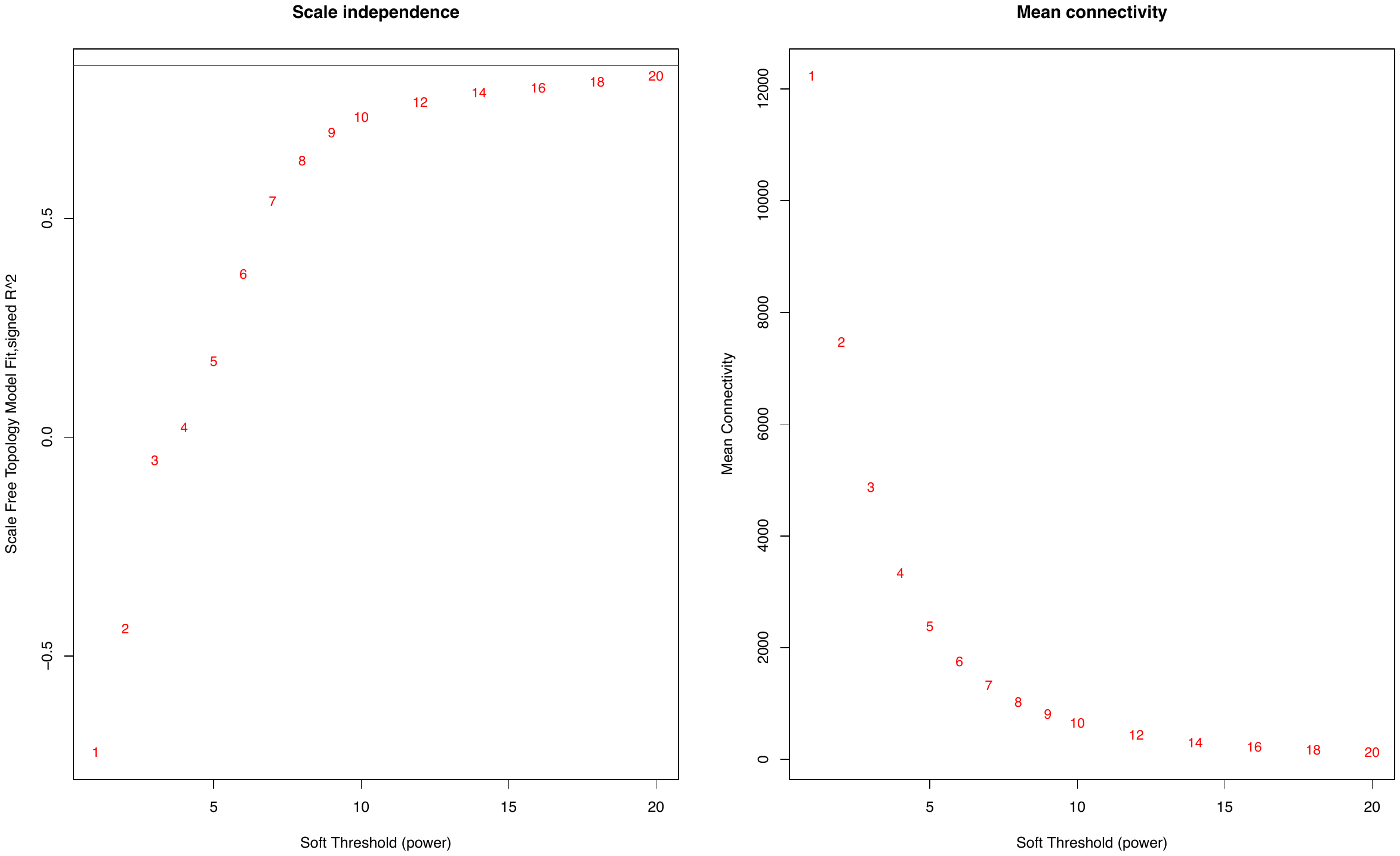
